# RNA triplex structures revealed by WAXS-driven MD simulations

**DOI:** 10.1101/2022.02.13.480274

**Authors:** Yen-Lin Chen, Weiwei He, Serdal Kirmizialtin, Lois Pollack

## Abstract

RNA triple helices are commonly observed tertiary motifs that are increasingly associated with critical biological functions, including signal transduction. Because the recognition of their biological importance is relatively recent, their full range of structures and function has not yet been elucidated. The integration of solution wide-angle X-ray scattering (WAXS) with data-driven molecular dynamics (MD) simulations, described here, provides a new way to capture the structures of major-groove RNA triplexes that evade crystallographic characterization. This method yields excellent agreement between measured and computed WAXS profiles, and allows for an atomically detailed visualization of these motifs. Using correlation maps, the relationship between well-defined features in the scattering profiles and real space characteristics of RNA molecules is easily defined, including the subtle conformational variations in the double-stranded RNA upon the incorporation of a third strand by base-triples. This readily applicable approach provides unique insight into some of the interactions that stabilize RNA tertiary structure and enable function.

## Introduction

Recognition of RNA’s importance as a powerful tool in supporting life and fighting disease continues to grow. An improved description of its structures is essential to further understand RNA’s regulation of biological function and to advance its application as a therapeutic^1^ and/or vaccine.^2^ Many RNAs contain base paired, duplex regions, connected by a variety of different motifs that often position unpaired nucleotide bases for additional stabilizing interactions. These latter interactions vary in type, they can involve long range base pairing, or other more intricate tertiary interactions that secure distal parts of the molecule. A survey of these interactions can be found in Ref.^3^

An intriguing class of RNA tertiary motifs contain base triples. Formed by stacked Hoogsteen base triples U-A·U and C-G·C^+^, RNA triplexes are now recognized as tertiary motifs^3–5^ in naturally occurring functional RNA architectures.^6^ They are found in human telomerase,^7–10^ spliceosomal RNAs,^11, 12^ long noncoding RNAs (lncRNA),^13–15^ viral RNAs,^16–19^ and riboswitches.^20–22^ Recent work underscores how triplex formation can transmit biological signals by sequestering the polyA tails of mRNA.^15^ Despite their potential biological or technological importance,^23^ the structures of only a few longer triplex structures have been solved (e.g. PDB id 6SVS^24^). Although these structures possess well defined contacts, the degree of flexibility displayed by the third, triplex forming oligomer, is unknown, yet is likely important in understanding how these molecules interact with partners. The goal of this work is to explore the conformation(s) of RNA triplexes that incorporate both stable and flexible regions, and to study the counterion distribution around such highly charged structures.

There has been much recent interest in developing accurate, atomically detailed models of RNAs. In contrast to well-ordered proteins, all atom MD simulations present challenges for RNA,^25^ including concerns about force field accuracy or sampling issues resulting from RNA’s rough energy landscape.^26^ Despite recent efforts ^27, 28^ it is not yet clear if current approaches provide a general framework to provide atomically detailed descriptions consistent with experiments. As an alternative approach, all atom models of RNAs are being successfully refined and tuned to come to agreement with experimental data acquired by NMR,^29–33^ cryoEM^34^, solution small^35^ or wide angle scattering (SWAXS).^36^

A recent study that motivates this one combines the Sample and Select (SaS) approach with wide angle x-ray scattering data on RNA duplexes.^36^ Results provide insight both into variations from A form structure and counterion distributions around double stranded RNA. To advance this work and to include higher order interactions, we made a similar comparison between SWAXS measurements of constructs containing multiple, stacked RNA base triples, and computed profiles of an ‘ideal’ triplex, subject to extended sampling. This SaS approach failed to capture the details reported in the experiment; the ‘correct’ structures do not appear to be present in the pool. This disagreement motivated the application of the data-driven, all atom MD simulations to enhance our previous SaS approach, described herein. This method relies on tight coupling of simulation with solution X-ray scattering data and is assisted by the undulations that appear in the higher angle solution scattering, reflecting the strong structural periodicities of the linked phosphate backbones. These features are used to ‘steer’ the MD simulation towards the experimental signal. Subtle variations of the structures are interpreted using correlation maps.^37^ As mentioned above, similar data-guided approaches are being exploited in conjunction with other experimental measurements ^29–35^.

This integrated approach reveals solution structures of hairpin RNAs alone and complexed with triplex forming oligomers (TFO’s). Of great interest are the changes to the duplex structure that accommodate and stabilize base-triple interactions. The atomically detailed structural ensembles afforded by MD simulations reveal changes to the cation atmosphere upon triplex formation, and resolve subtle differences in the measured WAXS signals that depend on the presence of particular (physiological) salt ions. Strikingly, our findings elucidate the unique role of U·A-U units in triple helical structures, offering biophysical insight into the interactions that stabilize RNA triplexes. Finally, although demonstrated in triplexes our method can be modified to elucidate the structural ensemble of more complex RNA molecules, as long as sufficient features are represented in experimental curves to serve as a guide.

## Materials and Methods

### RNA Triple Helix Motif Design

We designed short and long RNA hairpin constructs, called ST-DPX and LT-DPX, by terminating 17- and 29-base-paired (17- and 29-bp) duplex stems respectively with a UUCG tetraloop. The stems consist of sequence-specific 12-bp and 24-bp binding sites for two TFO’s, of 12 and 24 nucleotides in length, called TFO12 and TFO24, forming major-groove RNA triple helical structures, denoted ST-TPX and LT-TPX. The schematic secondary structures of the aforementioned RNA molecules are shown in Fig. 1(a). See section **RNA Triple Helix Motif Design** in ***SI Appendix*** for a detailed description.

**Figure 1:**
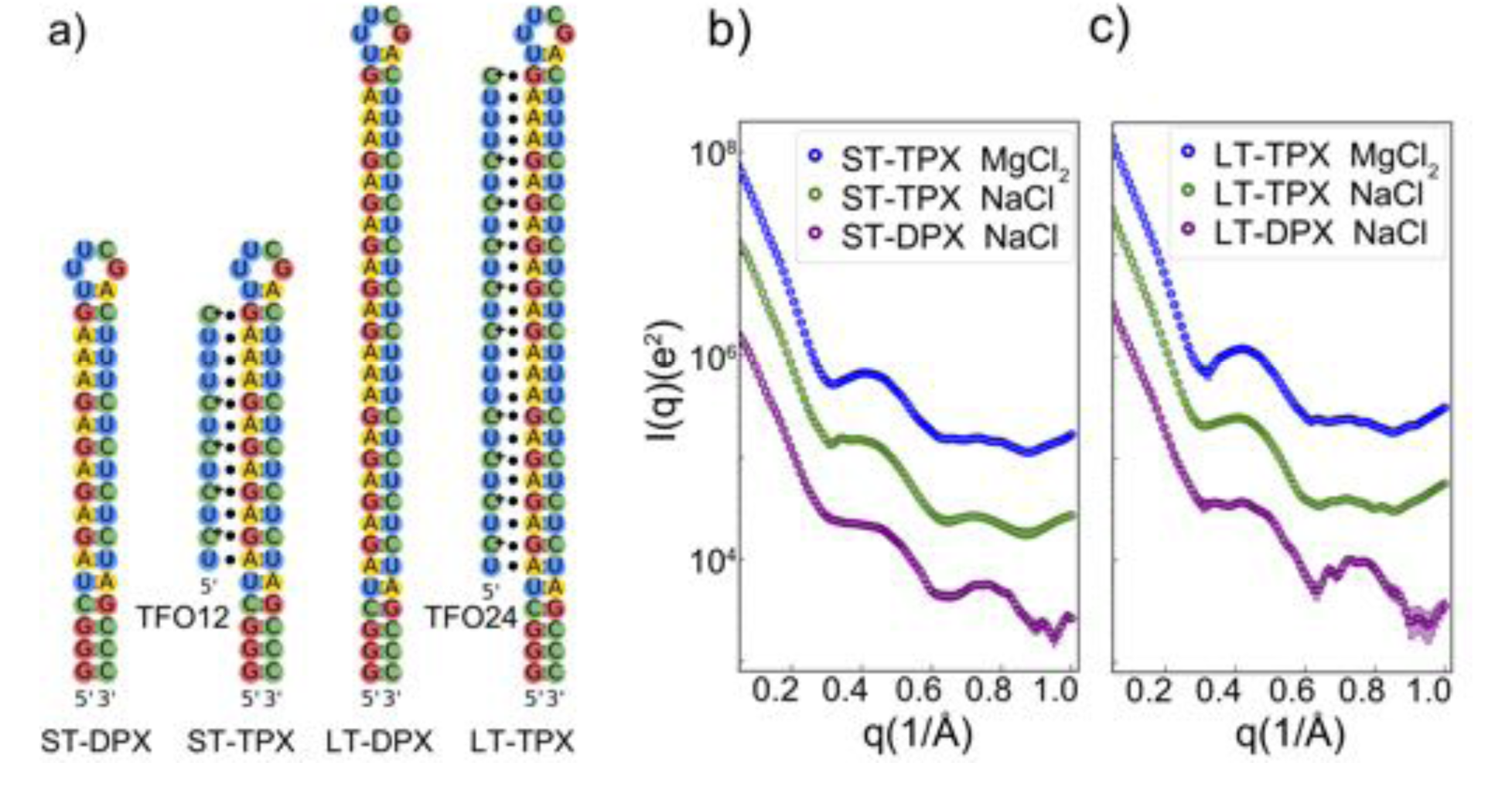
(a) Schematics showing the RNA duplex and triplex motifs designed for this work. The Short-Tetraloop Duplex (ST-DPX) has a UUCG tetraloop and a 17-base-paired stem with one TFO12 binding domain. A custom designed 12 nucleotide long TFO is added to yield multiple base triples, forming an RNA triplex: ST-TPX. We doubled the TFO12 binding domain to create a triplex forming partner for the Long-Tetraloop Duplex (LT-DPX). When bound, the duplex-TFO24 pair comprises the longer RNA triplex: LT-TPX. See main text for a more detailed description. (b-c) Solution X-ray scattering profiles of the different motifs and salts studied: ST-TPX/LT-TPX in 5 mM MgCl_2_ (blue), ST-TPX/LT-TPX in 200 mM NaCl (green) and ST-DPX/LT-DPX in 200 mM NaCl (purple).

### RNA Triple Helix Sample Preparation

Both ST-DPX and LT-DPX RNA molecules were synthesized using 4 batches of 10 T7 reactions at 37 °C from dsDNA templates purchased from IDT (Coralville, IA). We used ethanol precipitation at -80 °C to stop the T7 reactions and condense the RNA samples with impurities. The RNA samples were first purified using a Mono Q 5/50 GL anion exchange column (GE Healthcare, Chicago, IL) at pH 7.0 then annealed in buffers containing 40 mM NaCl, 20 mM sodium 3-(N-morpholino)propanesulfonic acid (Na-MOPS), 50 *µ*M ethylenediaminetetraacetic acid (EDTA), pH 7.0. Half of each ST-DPX and LT-DPX sample was then annealed with the protonated TFO12 and TFO24 strands in 200 mM NaCl, 1.0 mM MgCl_2_, 20 mM sodium 2-(N-morpholino)ethanesulfonic acid (Na-MES), 50 *µ*M EDTA, pH 5.5 to form ST-TPX and LT-TPX respectively. ST-TPX and LT-TPX were anion-exchange purified and buffer exchanged to either 200 mM NaCl or 5 mM MgCl_2_ for solution X-ray scattering experiments. A detailed description of sample preparation and characterization can be found in section **RNA Triple Helix Sample Preparation** of ***SI Appendix***.

### Solution X-ray Scattering

All solution X-ray scattering data were acquired at the 16-ID (LiX) beamline of the National Synchrotron Light Source II (NSLS II) at Brookhaven National Laboratory.^38^ Sixty *µ*L of solution samples and their matching buffers were manually loaded and individually measured in a continuous flow mode with five 2-second exposures. The momentum transfer of the scattered X-ray photons is defined as *q* = (4π/λ) sin θ, where λ is the incident X-ray wavelength in Å and θ is half of the scattering angle. Solution X-ray scattering data with a *q* range of 0.005 to 3.2Å*^-^*^1^ were recorded using a Pilatus 1M (small angle) and two Pilatus 300K (wide angle) detectors (Dectris, Switzerland) in vacuum. On-site frame-to-frame data screening was done by *py4xs* package^38^ to ensure matching of pre- and post-sample buffers and to ensure the absence of radiation damage. For all measurements, the signal-to-noise ratio (SNR) was above 30.0 at *q* = 1.0 Å*^-^*^1^. The SASBDB access codes for experimental profiles used in this study are reported in Table S2.

### Calculation of Solution X-ray Scattering Profiles

Small- and wide-angle X-ray scattering profiles were calculated following the scattering theory from excess electron densities.^39–41^ We used a probability isosurface to enclose the ion and solvent shells near the RNA molecule with a distance of *d* = 12 Å from the RNA surface. This distance was determined empirically. Fifty frames of solvent were used to estimate the electron density of the bulk solvent and to account for the excluded volumes. The orientational integral of the scattering amplitudes over 4π solid angle was computed numerically, using 1,750 uniformly distributed *q* vectors on a sphere,^42^ spanning a *q* range from 0.0 to 1.0 Å*^-^*^1^ with 201 points.

### The Customized Statistical Metric

The agreement between experimentally and computationally acquired solution X-ray scattering profiles was quantified using the customized 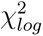 metric:

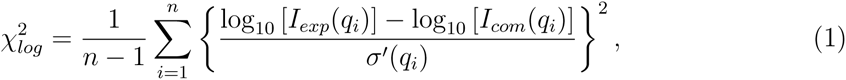

where *n* is the number of *q* points and *I_com_*(*q_i_*) and *I_exp_*(*q_i_*) are the computed and experimental intensities at *q_i_* respectively. In Eq. 1, cr*^0^*(*q_i_*) denotes the propagated error at *q_i_*: cr*^0^*(*q_i_*) = cr(*q_i_*)*/*[*I_exp_*(*q_i_*) log 10] with cr(*q_i_*) being the experimental error. 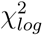 reduces the weight on the small-angle data and is statistically similar to linear x^2^ given the large signal-to-noise ratio (SNR *>* 10.0).^43^

### Molecular Modeling and Simulation Methods

Initial models of the ST-TPX, LT-TPX, ST-DPX, and LT-DPX were built using 3D-NuS,^44^ Nucleic Acid Builder (NAB)^45^ and 3dRNA^46^ programs. The constructed models were energy minimized using the GROMACS 5.0.5 package^47^ with the Amber xOL3 force field.^48, 49^ Subsequently, the constructs were solvated with water and ions to enable explicit water simulations. To mimic experiments, we prepared the RNAs in solutions containing ions at 200mM free Na^+^ and 5mM free Mg^2+^ concentrations. We used xOL3 parameters for nucleic acids, Smith and Dang parameters for ions ^50^, and modelled water using TIP3P.^51^ Details of the general simulation method are summarized in section **General MD simulation set up** of ***SI Appendix***.

### Sample and Select Approach

300 ns long room temperature (300K) production runs were employed to compute the initial WAXS profiles. Starting from the last frame of the MD simulation we performed a 100 ns-long high-temperature (HT, T=340K) simulation for each system. The sampled conformations are clustered based on their RMSD using the gromos algorithm.^52^ WAXS profiles were computed using the method described above for selected cluster centers. We evaluate the fit of each cluster using the metric defined in (Eq. 1). The cluster that gives rise to the smallest x^2^ is selected for the WAXS data-driven simulations.

### WAXS-driven MD simulations

The incorporation of experimental WAXS data [*I_exp_*(*q*)] to MD simulations was achieved by a Hamiltonian including a hybrid energy *E_hybrid_* = *E_FF_* + *E_W AXS_* term, where *E_FF_* is the energy from the MD force field, while *E_W AXS_* represents the energetic penalty if the real-time computed amplitude [*I_comp_*(*q, t*)] deviates from the target scattering curve [*I_exp_*(*q*)] of the experiment. The real-time WAXS curves are calculated on-the-fly from the simulation using the method described in **Calculation of Solution X-ray Scattering Profiles** and in Ref.^53^ The energy penalty to the deviation between the simulated and experimental WAXS profiles started with a logarithmic non-weighted coupling term,

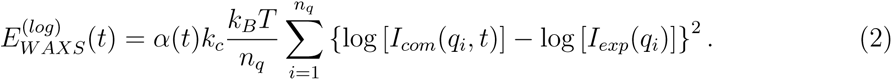

Here, *n_q_* is the number of intensity points spanning the specific range of scattering vectors *q*. The coefficient *k_c_* adjusts the weight of the WAXS potential *E_W AXS_* compared to the force field potential *E_FF_* . The parameter ff(*t*) is a time-dependent function that allows the gradual introduction of external coupling at the start of the simulation. Later, a nonweighted coupling was employed to reduce the curve deviation at higher *q* regions, followed by uncertainty-weighted coupling for globally fitting the WAXS signatures. Further details of the WAXS-driven MD approach can be found in section **Coupling experimental data with MD simulations** and **WAXS-driven simulation set up** of ***SI Appendix***

### Pairwise Distance Correlation Analysis

To evaluate the correlation between structural features and WAXS signals, we construct correlation maps (Eq.11 of ref^36^). We computed the normalized deviation between logarithmic intensities of simulated and experimental curves. To monitor structural changes, we used normalized deviation of pairwise distances. These distances were measured by picking the phosphate (P) atom of each residue, to which the solution X-ray scattering is most sensitive. The logarithmic form of the intensities allowed us to focus on structural characteristics beyond *d_min_* = 2π*/q_max_* ≈ 5Å. The normalized deviation at angle *q_i_* was computed by

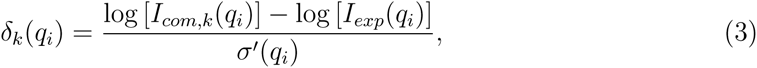

where, *I_com,k_* (*q_i_*) and *I_exp_*(*q_i_*) are intensities from computation and experiment respectively. Accordingly, the structural fluctuation of the *k*-th conformation was assessed using:

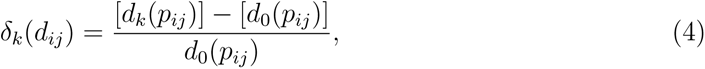

where, *d_k_* (*p_ij_* ) is the pairwise distance between residue *i* and residue *j* of the *k*-th conformation, and *d*_0_(*p_ij_* ) is the equilibrium value of that distance.

### Data Analysis

The conformations which display 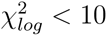 (Eq. 1) were collected and used for all subsequent analysis. To investigate Loop*<->*Duplex and TFO*<->*Duplex positioning, we carried out contact analysis (See **Residue-residue Contact Analysis** in ***SI Appendix*** ). To study duplex topology, we estimated the groove width using the 3-DNA program.^54^ The structure of the cation cloud around the helices was analyzed using:

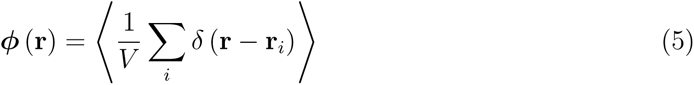

where **r** ≡ (*x, y, z*), describes an arbitrary position in Cartesian coordinates, *ϕ*(*x, y, z*) is the average cation number density, *V* is the volume of simulation box, c(*x*) is the Kronecker delta. The sum is performed over the cation index *i*, and 〈…〉 represents the ensemble average.

## Results

### WAXS reveals structural features of RNA Triplex

Solution X-ray scattering provides both global and local information about the structure(s) of RNA. While small-angle X-ray scattering (SAXS) reveals the overall size and shape of the molecule, wide-angle X-ray scattering (WAXS) is now a proven tool for discerning variations in the solution structure(s) of RNA molecules,^36, 55, 56^ importantly reporting higher-resolution information about the distribution of distances present in the molecule *in vitro*. We apply WAXS to understand the structure(s) of motifs with tertiary structures: triple stranded RNAs.

To obtain triplex structures for measurement, we designed two duplexes as well as a complementary single-stranded TFO that binds to each duplex (see **RNA Triple Helix Motif Design** and **Sample Preparation**). We subsequently refer to the duplex motifs as ST-DPX, and LT-DPX for short and long constructs. Their respective triplex counterparts are referred to as ST-TPX and LT-TPX.

The secondary structures of the molecules studied here are shown in Fig. 1a. Figure 1b-c shows their WAXS profiles. Profiles of the shorter duplex and triplex (ST-DPX and ST-TPX) are shown in the left panel, in solutions containing different salt ions. Profiles of the longer constructs are shown in the right panel. The difference between duplex and triplex scattering is evident from the curves. The peak in the triplex scattering profile (near *q*=0.4 Å^−1^) flattens for the duplex and the peak in the duplex scattering profile (near *q*=0.75 Å^−1^) flattens for the triplex. The WAXS profile of the triplex sample is sensitive to the surrounding ions; features in divalent Mg are better defined than those in monovalent Na (Note: a similar effect was observed for duplexes, as reported in^36^). Finally, comparison of panels b to c in Fig. 1 shows that the WAXS features are better defined in the longer constructs. This is not surprising given the increased reinforcement of correlations with length in a regular structure.

### Computational method generates WAXS consistent RNA ensembles

The sensitivity of WAXS to the variations with length, sequence and salt is promising; however, the information hidden in WAXS curves demands computational modelling to provide accurate insights. To reveal the structural differences implied by WAXS, we employed a computational strategy that is summarized in Fig. 2a. Simulations begin with an RNA model, built from only secondary structure information (Fig. 1). We use this model structure to generate an ensemble of conformations using MD simulations that account for the RNA flexibility, explicit water and cations (Fig. 2b). Details of our MD methodology can be found in methods and SI (see **WAXS-driven MD simulations**). We compute the WAXS profiles from the atomic coordinates. Later, we assess the agreement of the profile derived from computational pool by comparing it to the measured scattering profiles from experiment, using a customized x^2^ metric (see **The Customized Statistical Metric**). If agreement is not achieved, a high-temperature (HT) MD simulation is used to expand the structural pool. The HT trajectory is clustered to identify important conformational sub-states. Each cluster is then evaluated against experimental data. If the x^2^ of the best cluster is satisfactory we select the conformation and further sample conformations around it using brute force MD simulations. As shown in our past work,^36^ this SaS strategy can be effective if the issue is sampling, meaning that the ’correct’ structures are present in the pool, but may be difficult to access, given the starting point of the simulation (A-form duplex). For more complex structures, where the ‘correct’ models may not be present within the computational pool, a new approach is required to refine the model. The WAXS-driven MD simulations, described here, allow us to steer the best conformation of the SaS simulation in new directions to achieve the requisite agreement. After the starting conformation is selected, our WAXS-driven approach guides the MD towards regions of conformational space where the computed profiles find better agreement with the experimental data. Fig. 3a, (black) shows results for the ST-TPX RNA in NaCl. The process, as applied to the other RNAs studied, yields similar improvements in fit and is reported in Fig. S4. The structural search reaches our set criteria (x^2^ *<* 2) within 40 ns (Fig. 3b). Once the search converges to a structure consistent with experimental signal the error stays around the same value suggesting no major change in WAXS profiles. To provide ensembles for analysis we construct a pool from the conformations that fall below the threshold. These pools in each condition are used to derive equilibrium properties and distributions. Throughout the course of the conformational search stage of the simulation, the error and the structural changes are simultaneously monitored (Fig. 3b & c). Interestingly, significant deviations from the starting structure are localized to specific regions, for example near residues 19-22 and 41-52 in ST-TPX. Changes are ’tracked’ by monitoring structural deviations in these regions, which correspond to the tetraloop and TFO position of the ST-TPX respectively. As a control, we employ unbiased MD starting from the initial RNA structure and from the best structure selected by WAXS-driven MD (Fig. 3a, red versus cyan green). The agreement of unbiased MD is poor in both cases, highlighting the necessity of our approach. Specifically, the spontaneous drift of the structural pool from the best structure suggests discrepancies in the forcefield in their representation of triple-helix RNAs. In contrast, computed WAXS profiles of duplex structures evolve towards better agreement with experiments when sampling starts from the right conformational region (Fig. S4 a,c). Thus our current forcefield appears to adequately capture duplex topology, but needs adjustments when modeling higher order structures.

**Figure 2:**
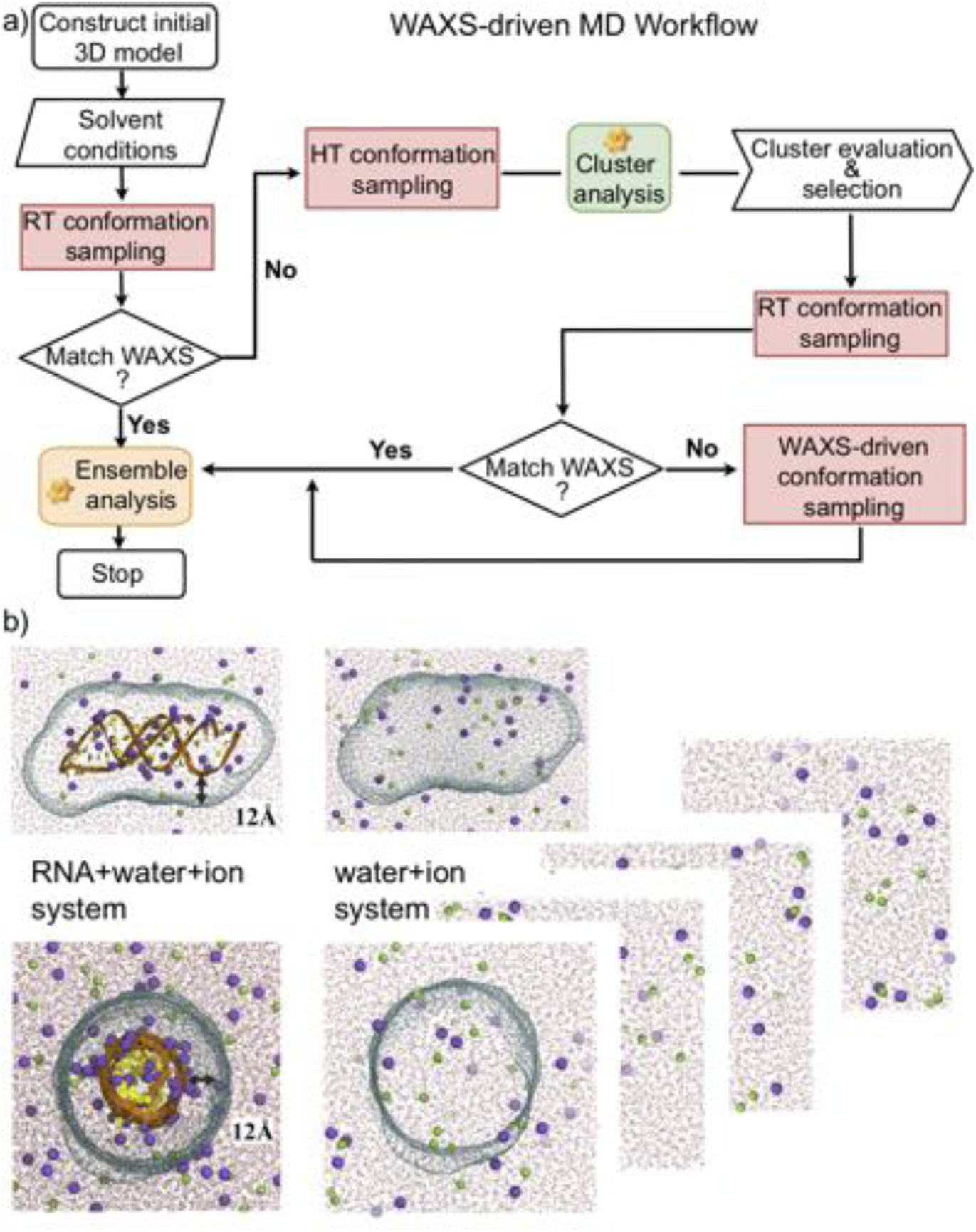
(a) The flowchart of WAXS data-driven MD approach. Two feedback loops are included, and the path through is determined by the agreement (or lack of agreement) with WAXS profiles. If agreement is not achieved in the RT pool, the SaS approach^36^ is incorporated to enhance the sampling. The second feedback loop is triggered when the customized metric in Eq. 1 of all the sampled conformations exceeds the threshold. After multiple iterations, the data-driven strategy generates a conformation pool satisfying the experimental WAXS data, assessed by the value of our customized metric. (b) Illustration of the RNA+water+ion system and water+ion system used in the WAXS data-driven MD simulations. The molecular envelope constructed from a 3D probability density isosurface 12Å from the helix surface to encompass the solvent/ion shell is depicted by mesh. The scattering intensity of water+ion system was computed by applying the same envelope construct.

**Figure 3:**
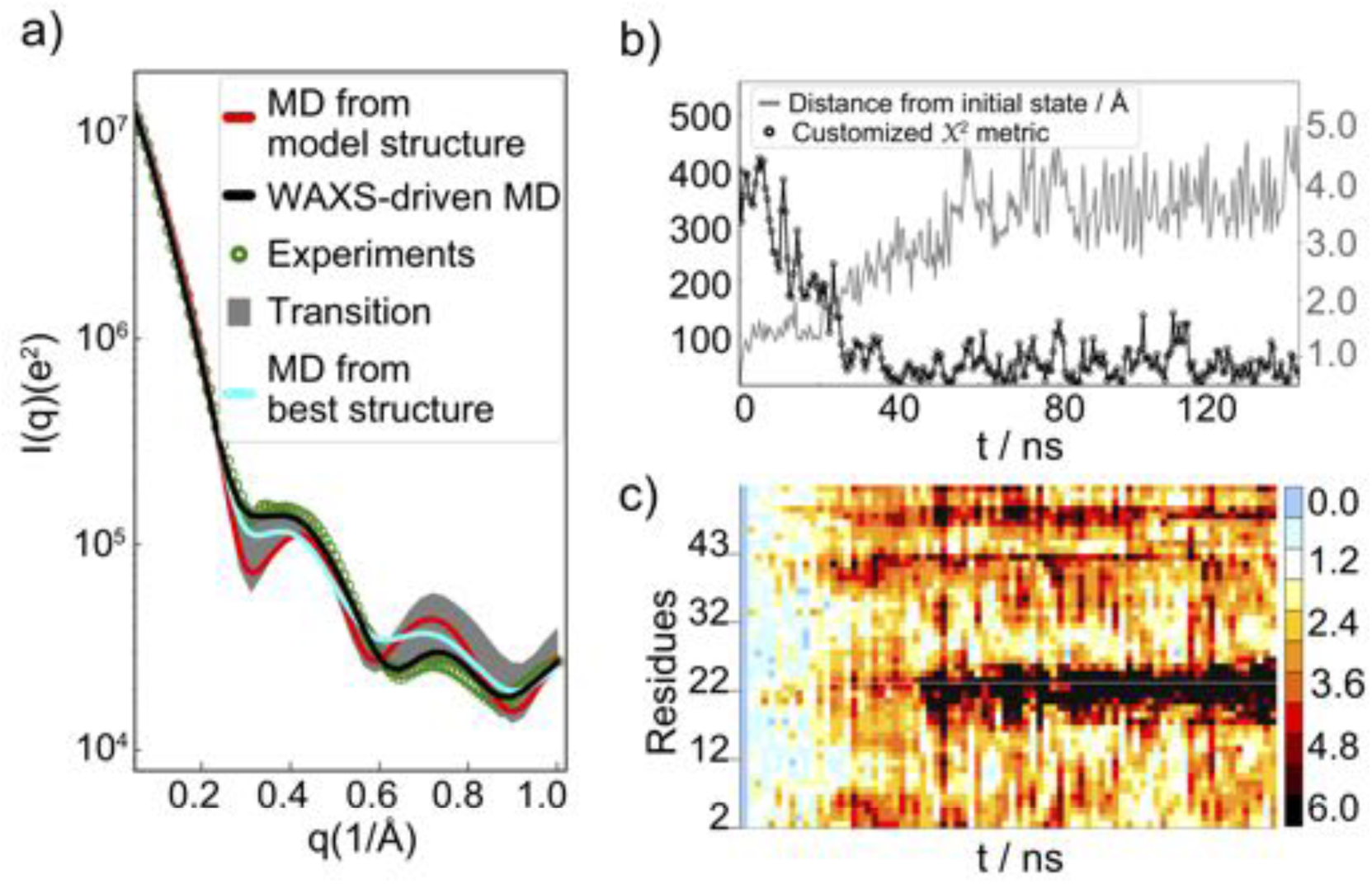
Illustration of WAXS data-driven approach based on simulation of ST-TPX in buffered solution containing 200 mM NaCl. a) The experimental data that are the target of the WAXS-driven MD simulations are shown as green circles. The scattering profile computed from the starting MD structure is shown in red, the profile computed from the best-fit MD structure is shown in cyan, the progression of scattering profiles during the data-driven simulation is shown in gray, denoted as transition in the figure and the result of the data-driven simulation is shown in black. The WAXS evolution of the remaining constructs is shown in Fig. S4. b) The time variation of x^2^, as well as the overall RMSD (Å) change are shown, using the initial frame as reference. The asymptotic amplitude and turning point of the two variables respectively determine the convergence of simulations. c) The time evolution of the RMSD at the level of individual residues is shown, using the initial structure as the reference. The color bar provides the distance from initial conformation. Note the relatively higher structural deviations displayed near residues 19-22 and 41-52 residues which correspond to the tetraloop and TFO of ST-TPX, respectively.

### Pairwise distances correlates with WAXS profiles

Key to the success of this method is understanding how WAXS features correlate with RNA structural features. Here, we benefit from data science. Structures sampled by MD simulation serve as a training set to find relationships between features in the computed WAXS profile and 3D structural details. We first examined the correlation between residue pair distances *d_ij_* and WAXS amplitude at each discrete point *q_k_* , *I*(*q_k_* ). (See detailed description in **Pairwise Distance Correlation Analysis**). Correlation maps are used to display these relationships. Fig. 4 summarizes our analysis for ST-TPX. A comparable analysis for LT-TPX is detailed in Fig. S8.

**Figure 4:**
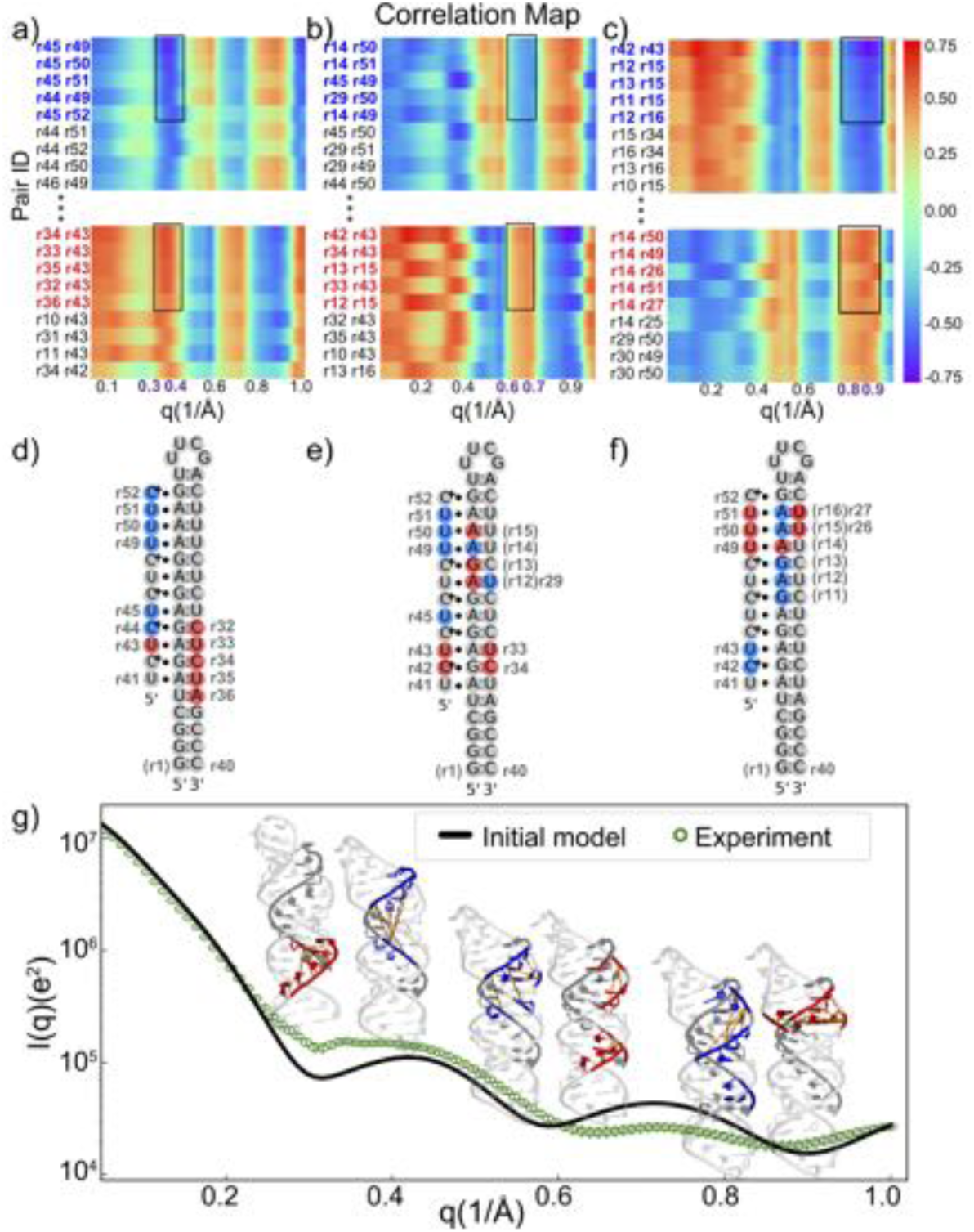
Correlation analysis based on comparison of measured WAXS profiles with structures and profiles computed from standard MD simulations of ST-TPX in 200 mM NaCl. a-c) These partial correlation maps, extracted from the full correlation map (Fig. S6), display the distance between pairs of phosphate atoms associated with the indicated residues, computed from MD structures, against the normalized deviation of the computed WAXS profile from the experimental measurement. These maps identify structural regions that potentially contribute to deviations in specific *q* regions. Regions with the highest deviations in a given *q* range, e.g. the residue pairs with high correlations (*|ρ| >* 0.5 & 5 largest values), are highlighted within black rectangles. We focus here on structural regions that create significant deviations in profiles near *q* ≈ 0.35Å*^-^* (a), *q* ≈ 0.65Å*^-^*^1^ (b) and *q* ≈ 0.85Å*^-^*^1^ (c). d-f) The regions identified in parts (a-c) are mapped onto the secondary structure and real space models of the RNAs. Residue pairs that contribute to changing the scattering profile near *q* ≈ 0.35Å*^-^* (d), *q* ≈ 0.65Å*^-^*^1^ (e) and *q* ≈ 0.85Å*^-^*^1^ (f), respectively are highlighted in color. Panel (g) connects distinct structural variation with changes in the scattering profiles. A comparable analysis applied to LT-TPX is provided in Fig. S8.

The normalized pairwise distances of Fig. 4 show both positive and negative correlations with structural features. A positive correlation implies an increase in *I*(*q*) with an increase in normalized deviation of distance pair at a particular *q* value, with the opposite true for negative correlations. Here, we show only the highest correlations in magnitude for three *q* ranges (0.3-0.4 Å^−1^), (0.6-0.75 Å^−1^), and (0.8-0.95 Å^−1^) (Fig. 4).

These analyses demonstrate that the TFO conformations play a significant role in shaping the scattering profiles (Fig. 4). As an example, the experimental profile deviates significantly from the unbiased MD prediction near *q* = 0.35 Å^−1^. The correlation map shown in Figure 4a provides insight into the structural changes that can bring the experimental and predicted curve into closer agreement at this *q* value. Specifically, a decrease in distance between pairs highlighted with blue blocks, as well as a flipping out of the end of the TFO, which increases the pairwise distances shown by the red blocks, suffice to increase the computed scattering intensity at this *q* value, smearing the obvious minimum that is present in the standard MD curve. This change corresponds to a compression of the TFO along its length (decreased distance between the blue-colored residues in Figure 4d), as well as an increase in the distance between its 5’ end and the duplex. Together, these effects enhance the signal intensity at *q* = 0.35 Å^−1^ to bring the computed and experimental curves into agreement (Fig. 4a,d,g). Similarly, the discrepancy between unbiased MD and measurement seen at around *q* = 0.65 Å^−1^ can be compensated by breaking the symmetry in the duplex-TFO pairwise distance (Fig. 4b,e). This analysis suggests that variations in the structure may not always be reflected at a unique *q* range, rather, the auto-correlations among WAXS features (see Fig. S5a for ST-TPX) suggests that structural variations may induce changes at multiple *q* regions. Despite the complexity that this multiple dependency can induce, these cross correlations do insure consistency. For example, the WAXS features at around *q* ≈ 0.35 Å^−1^ positively correlate with that of *q* ≈ 0.65 Å^−1^, however the same feature negatively correlates with intensity at *q* ≈ 0.85 A^\x{FFFF}^*^-^* (Fig. S5a). Therefore, a closer duplex-TFO contact amplifies the WAXS amplitude at *q* ≈ 0.65 A^\x{FFFF}^*^-^* (Fig. 4b,e,g, blue blocks) and at the same time, reduces the *I*(*q*) at *q* ≈ 0.85 A^\x{FFFF}^*^-^* (Fig. 4c,f,g, red blocks). Meanwhile, stem expansion near the loop increases the amplitude at *q* ≈ 0.85 A^\x{FFFF}^*^-^* (Fig. 4c,f,g). Thus, the agreement of our computed profile with experiment over the entire *q* range provides support for all of the selected changes.

The above discussion highlights our experimental sensitivity to the conformational state of the TFO. Additional information about changes in the duplex topology that result from TFO binding are also available: features in the WAXS profile correlate with the major groove width. The positive correlation at *q ⇡* (0.8-0.95 A^\x{FFFF}^ ), especially residue pairs of r14-r26 and r14-r27 (Fig. 4c,f), suggests a major groove widening which will be detailed in the next section. Despite the above-described success in articulating the structures of the duplex and the TFO, the correlation maps do not provide robust guidance (high values of correlation) for the changes in the tetraloop region (Fig. S6). This is not surprising, due the highly dynamic nature of the loops revealed by MD simulations,^57^ as well as the short length scales that characterize interactions within the loops. Data acquired at higher *q* values, beyond the scope of the present study, will be required to inform about the structure(s) of this region. The most representative structures from the WAXS-driven simulation are shown in panels b,c,d of Fig. 5.

**Figure 5:**
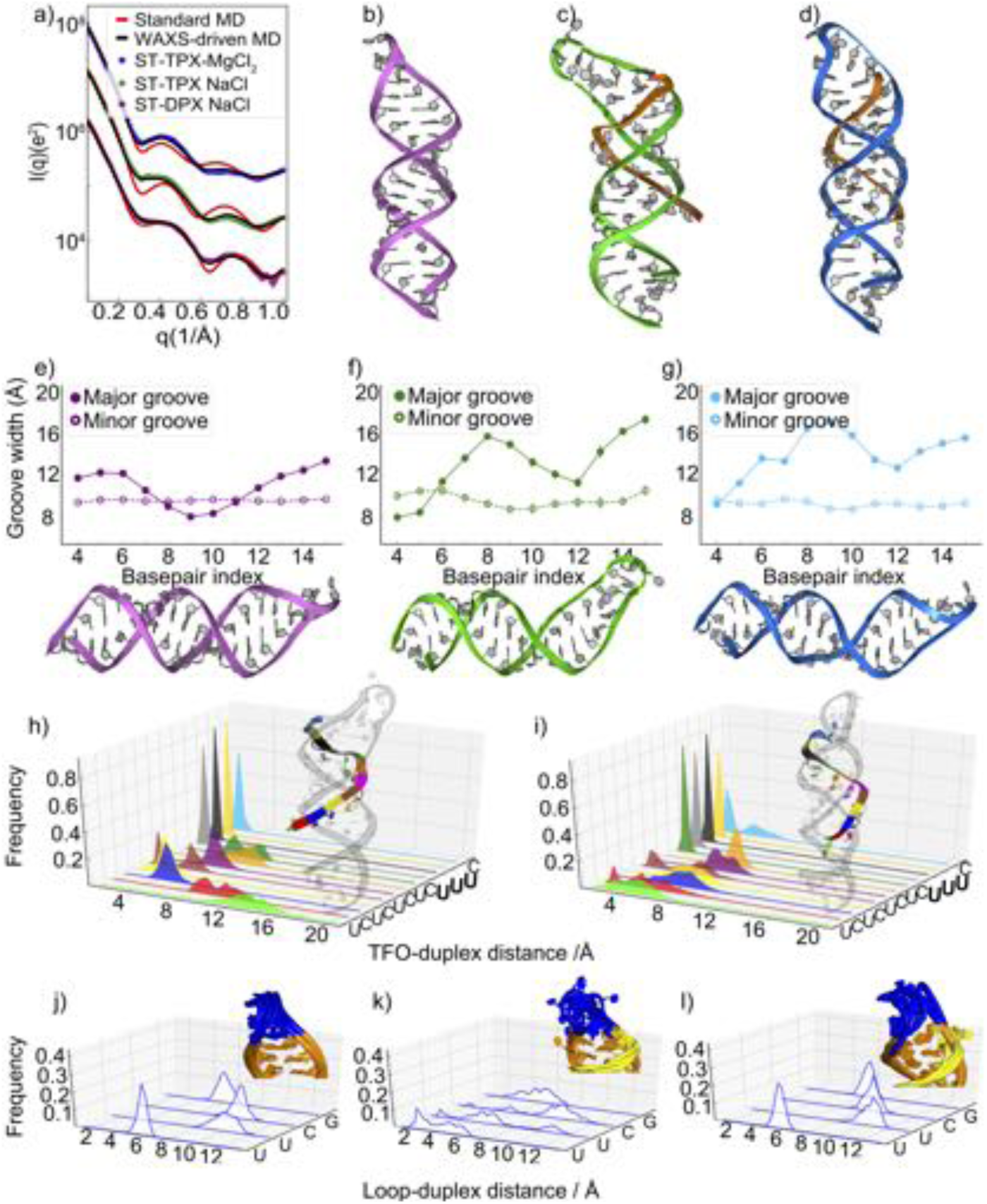
Structural analysis and comparison of short helices. (a) The experimental scattering profiles (circles), computed WAXS profiles from standard MD simulations of model structure (red) and WAXS-driven simulations (black) are pictured for three constructs. From the conformational ensembles generated by WAXS-driven MD, we show the most representative structure of (b) ST-DPX in NaCl, (c) ST-TPX in NaCl and (d) ST-TPX in MgCl_2_.For equilibrium properties we consider the whole ensemble. The ensemble was generated from conformations that fall below the threshold x^2^ *<* 2 value. (e-g) Average groove width (GW) of the duplex part of structures reveals a dependence on salt: (e) ST-DPX in NaCl, (f) ST- TPX in NaCl and (g) ST-TPX in MgCl_2_, respectively. (h-i) The contact analysis measuring the relative positions of the TFO and duplex of ST-TPX reveals detailed structural information about TFO binding in (h) NaCl and (i) MgCl_2_. (j-l) The contact analysis between tetraloop and duplex of (j) ST-DPX in NaCl and ST-TPX in (k) NaCl and (l) MgCl_2_ reveals an unexpected salt dependence of the loop within triplex structures.

### Duplex and triplex structures and how the TFO binding modulates RNA structural ensemble

We now turn to a detailed discussion of the structures of the ST-DPX and ST-TPX constructs which is uniquely enabled by the agreement of computed scattering profiles with experimental measurements. First, we note that unbiased MD simulations require only small modifications to accurately recapitulate the structure of the ST-DPX construct (Fig. 5a). In contrast, structural pools from unbiased MD simulations show much more deviation when modeling the triple stranded constructs (Fig. 5b-d). In these latter constructs, the final, MD-driven result, shown as structures in panels c and d, reveal significant variations in the tetraloop and TFO, which may be difficult for unbiased MD to capture. Once the simulations have been driven to agree with the experiment, the real space projection of the WAXS profiles afforded by all atom MD simulations provide a platform to investigate the TFO-induced structural changes with atomic detail.

To accurately describe and interpret the structural changes, we separately consider the three distinct components of the RNA structure: i) base-paired duplex stem (without TFO and loop), ii) TFO, and iii) tetraloop regions.

As a first step in providing a description of the duplex structure, as well as the ability to easily describe changes resulting from TFO binding, we focus on the dimensions of the groove widths (GWs). Figure 5e-g displays the widths of both the minor and major grooves for three short constructs. Figure 5e reports the average groove widths computed from the pool for the ST-DPX construct, revealing a consistently sized minor groove and some variations in the major groove width, especially near the ends of the base paired regions. Figures 5f and g show the significant TFO-induced changes to the major groove dimensions and therefore to the duplex structure. Most significantly, the major groove widens once the TFO is bound. This change is dramatic near the middle of the duplex and near the tetraloop (beyond basepair 14).

We now focus on the positioning and conformations of the TFO in the pool for the triplex construct in different salt conditions. To quantify our findings, we measured and report the pair distances between TFO and duplex nucleotides (details in method section **Residue-residue Contact Analysis**). These distance distributions allow for a comparison of sequence specificity of TFO binding across salt conditions and reflect the structural heterogeneity of Hoogsteen base triples. Most interestingly, the U*·*A*·*U base triples (UUU end, at the ’top’ in the figure) form close contacts to the duplex (Fig. 5h-i). This tight binding shows no variation with the salts used, suggesting a short range non-electrostatic force governing contact formation in these tandem uracil units. This observation confirms earlier work and crystallographic studies.^58–60^ While this ’upper end’ of the TFO appears to be fixed by the tandem uracil units, the contacts at TFO’s lower end appear more dynamic. This effect is more pronounced for RNA in Na^+^ in comparison to Mg^2+^ salt, and likely reflects the weakened electrostatic screening afforded by monovalent relative to divalent ions.

Finally, we consider the structure(s) of the loop region in our best-fit ensembles. As seen in the TFO contacts, a significant salt variation of this region is observed (Fig. 5j-l). Here the dynamics are depicted both as structures and as computed distances. Both representations illustrate the dynamic features of this flexible region. It is interesting to compare the loop structure in the ST-DPX and the ST-TPX complexes. Relative to the former, the loop structure in the latter tends to twist upon the introduction of the TFO (Fig. 5j,k). In addition, the ST-TPX loop appears more dynamic in Na^+^ than in Mg^2+^ (Fig. 5k,l). The added strand increases the charge density around the loop which likely leads to stronger repulsion that partially melts this region. With higher valence Mg^2+^, the canonical form of the tetraloop is restored (Fig. 5 l).

All together, we reveal that triplex formation requires global modifications of the duplex geometry, in both salt conditions studied. Salt dependent effects are manifested in the structural variations of the terminating loop as well as in the variations of structure of the TFO, especially near its lower end.

### Short and long triplex sequences show similar structural variations

To establish the generality of our findings, we measured and simulated systems consisting of longer, loop terminated, duplexes, with and without TFOs. We examined a longer stem-loop construct (LT-DPX), without and with the addition of a longer TFO (LT-TPX). The triplex forming constructs were examined in solutions containing either Na or Mg ions. Results for the LT-TPX show that the key deviations of MD predictions from WAXS measurements occur in regions that reflect structural coupling between the duplex and the TFO, consistent with findings from the ST-TPX system (Fig. S8). As in our above-described studies with the shorter motifs, we applied correlation analysis to bring the predictions for all of the longer constructs into agreement with experiment. Following the application of WAXS-driven MD, the representative best-fit structures of each case are shown in Fig. 6b-d. Conclusions from simulations of the structures of LT-DPX (Fig. 6e) are comparable to those derived from the ST-DPX system (Fig. 5e). In particular, increases in the major groove width are observed near the two ends of the helical regions, relative to near the center of the duplex. The minor groove widths remain relatively constant and unchanged from the shorter construct. For these longer constructs, the addition of the TFO induces a major groove widening, as seen in the shorter strands; however, it is more emphasized in LT-TPX where a nearly monotonic increase from bottom of the stem to the tetraloop is observed (Fig. 6f,g).

**Figure 6:**
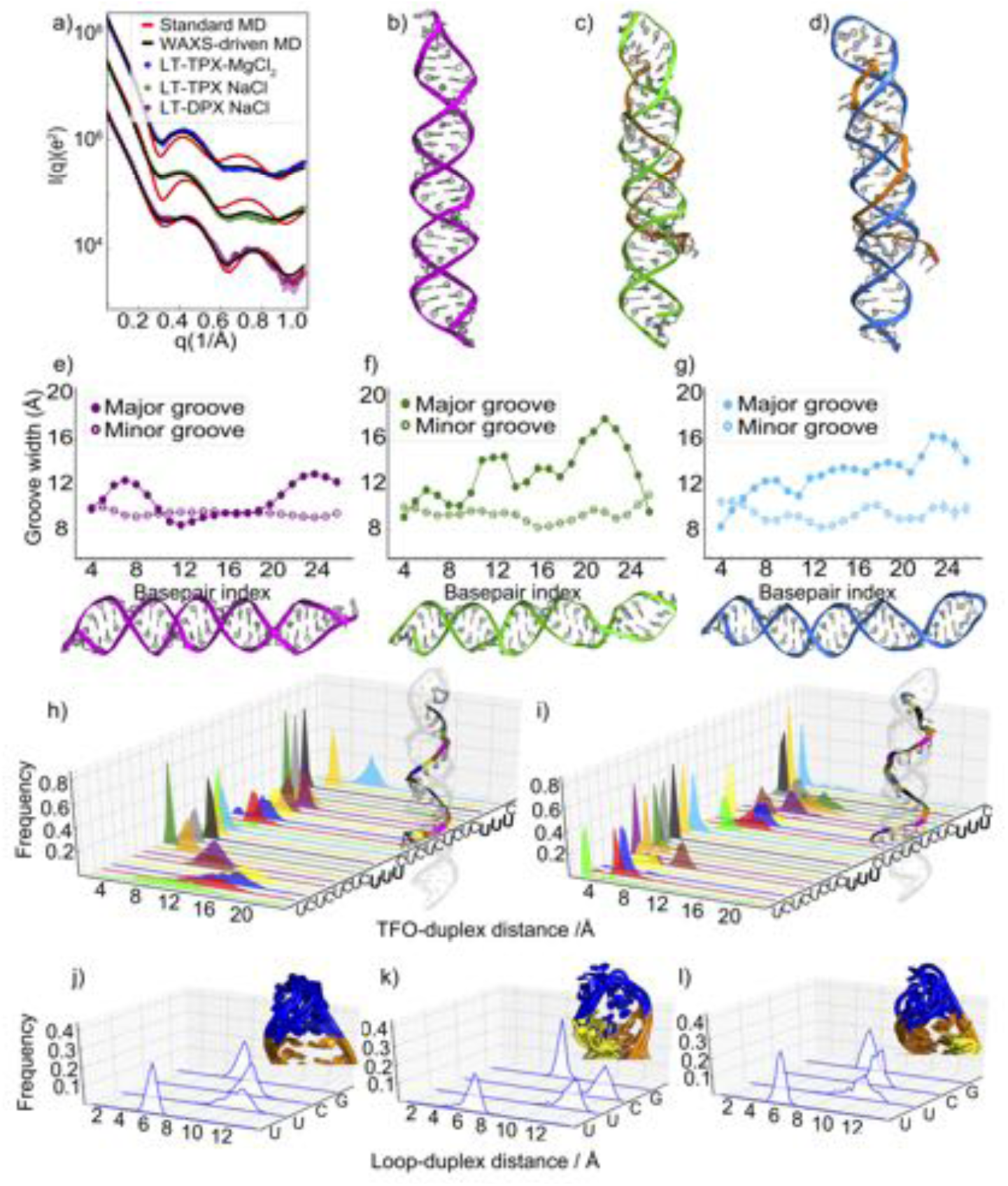
Structural analysis and comparison of constructs with longer helices. (a) The experimental scattering profiles (circle), computed WAXS profiles from standard MD simulations of model structure (red) and WAXS-driven simulations (black) are pictured for these longer constructs. As in Fig. 5, we show the most representative structure of (b) LT-DPX in NaCl and LT-TPX in (c) NaCl and (d) MgCl_2_. For equilibrium properties we consider the whole ensemble. The ensemble was generated from conformations that fall below the threshold x^2^ *<* 2 value. (e-g) Average groove width (GW) of the duplex part of structures in different salt conditions reveals a more distinct pattern than observed in the shorter complex for: (e) LT-DPX in NaCl, (f) LT-TPX in NaCl and (g) LT-TPX in MgCl_2_, respectively. (h-i) The contact analysis between TFO and duplex of LT-TPX in (h) NaCl and (i) MgCl_2_ and (j-l) between tetra-loop and duplex of (j) LT-DPX in NaCl and LT-TPX in (k) NaCl and (l) MgCl_2_ show smaller variations relative to the shorter construct in NaCl.

The binding mode of the longer TFO shows sequence specificity and salt dependence. Consistent with observations on the shorter triplex, the uracil bases of this TFO bind tightly to the duplex. The two tandem units of uracil share this trend (Fig. 6h,i) and the deeply buried major groove binding of these tandem units is salt independent. Due to its longer length, the end of this TFO (perspective view along y axis) is more susceptible to excursions and shows higher dynamical fluctuations, especially in Na salt. However, in these longer constructs the tetraloop exhibits less heterogeneity in NaCl (Fig. 6k) when compared to STTPX. Finally, in parallel with the ST-TPX, the structural heterogenity of the TFO terminus is restricted in Mg^2+^ (Fig. 6l).

### Spatial cation distributions of duplex and triplex structure are different

A great benefit of these atomically detailed, MD simulations is the additional information gleaned about how the spatial distributions of cations respond to accommodate TFO binding. These studies directly address the question of how a negatively charged third strand displaces tightly-bound cations from a deep and negatively charged groove. Fig. 7 shows the MD-derived average cation occupancy around the RNA structures in all of the systems studied.

**Figure 7:**
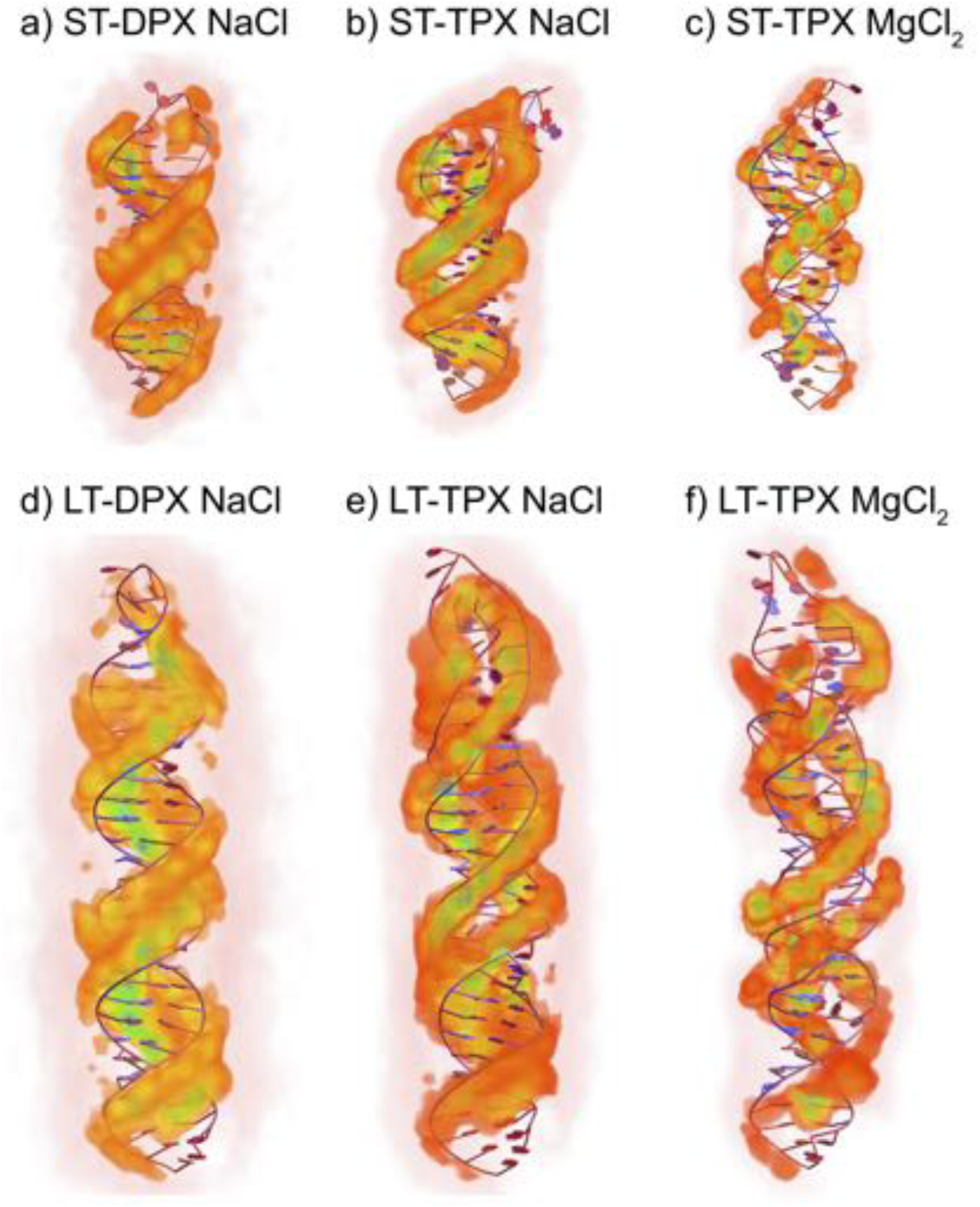
Three dimensional ion density profiles around the six construct studied, (a) ST- DPX in NaCl, ST-TPX in (b) NaCl and (c) MgCl_2_, (d) LT-DPX in NaCl, LT-TPX in (e) NaCl and (f) MgCl_2_, respectively. Increasing ion density is indicated with color: white(low) to red to green (high).

Marked differences are observed between cation distributions around duplex and triplex RNAs. This information is most relevant when the same salt is used, in this case NaCl. Consistent with our earlier works^61, 62^, duplex geometries show major groove binding of cations, (Fig. 7a,d), with localization of Na^+^ cations to the phosphate backbone. Most surprisingly, the binding of TFO to the major groove does not exclude Na^+^ cations from the major groove, rather, it leads to stronger localization of cations, into upper and lower layers juxtaposed to the third phosphate backbones (Fig. 7b,e). The tetraloop region also showed an increase in cation occupancy in the case of triplex structure in the stem region. This increased localization may result from the higher charge density created by the third strand. Similarly, hexa-hydrated Mg^2+^ cations also bind to the major groove of TPX structures despite the presence of TFO. Localization of divalent ions is more discrete (Fig. 7c,f), in sharp contrast to the case of Na^+^ where binding is territorial and diffusive.

## Discussion

Combining atomically detailed simulations with real experimental data can be a challenging task. Many factors impact the accuracy of the interpretation of the results, ^63–66^ such as the resolution of the experimental data, the accurate reflection in the measurement of well-defined structural correlations in the molecules, the type and magnitude of the errors, and finally the accuracy of the back-calculation.^67^ Computational challenges are also important, especially selection of a good starting model, assessment of overlap between the experimental and computational ensemble(s), and the role of forcefield and coupling methods on the ensemble generated need to be investigated more in depth to develop a robust computational strategy. Here, we proposed and demonstrated a strategy for solving the solution structure of RNA molecules possessing tertiary motifs. This framework integrates Sample and Select MD with WAXS-driven forces to direct the simulation into agreement with the experimental data. We applied this framework to solve the structures of biologically important, but structurally uncharacterized RNA triplex motifs. Structural variations, dynamic arrangements, as well as ion effects were accessible to our analysis, and were examined in depth. From these straightforward comparisons, we derived an improved understanding of RNA triplexes at the atomic level. For example, we observed that a like-charged third strand of RNA is readily accommodated by an already formed duplex, by widening the major groove, sequence-dependent contact formation and redistribution and localization of charge compensating cations. The new charge distribution pattern identified here, in addition to the structural importance of tight U.A-U base triples from tandem uracil residues likely contribute to function. They may contribute to long range molecular recognition and signal transduction in currently unrecognized ways. Our coupled studies also underscore the importance of divalent ions in stabilizing flexible motifs, tightening loops and lashing down free ends. These new degrees of freedom inform our understanding of RNA tertiary structure and inspire ongoing studies to more fully characterize the detailed electrostatic environment of these highly charged molecules, which uniquely enables their interactions with select cellular partners.

Our method fills an important void by integrating state-of-the-art wide angle solution X-ray scattering experiments with MD simulations to investigate ensembles of RNA structures. Though demonstrated for RNA triplexes, this general approach has the potential to solve a myriad of RNA structural motifs. Future applications will extend to study dynamic systems containing both protein and RNA partners.

## Acknowledgement

This work was supported by NIH Grant R35-GM122514 to L.P.. Support for work performed at the CBMS beam line LIX (16ID) at NSLS-II is provided by NIH - P30 GM133893, S10 OD012331 and BER- BO 070. NSLS-II is supported by DOE, BES-FWP-PS001. Computational research was carried out on the High Performance Computing resources at New York University Abu Dhabi. S.K. and W.H. are supported by AD181 faculty research grant. The authors thank Shirish Chodankar and Lin Yang for experimental assistance and Jorge Naranjo for his computational support. Author contributions: Y.-L. C. performed the experiments and analyzed the data. W.H. performed and analyzed the simulations. S.K. and L.P designed the study. All authors contributed to integrating experimental with computational data and to writing the manuscript.

## Supplementary Information

### RNA Triple Helix Motif Design

We designed an RNA hairpin construct called ”Short-Tetraloop Duplex” (ST-DPX), consisting of a UUCG tetraloop and a duplex stem with 17 base pairs:^23, 68, 69^ 5’- GGG CUA GAG AGA GAA AGU UUC GAC UUU CUC UCU CUA GCC C. The duplex stem further contains a 12-base-paired TFO12 binding domain, designed to form 12 consecutively stacked base triples, followed by 4 GC base pairs at the closing (blunt) end. The 12-base-paired TFO binding domain is an asymmetric sequence to ensure a specific binding direction of the partner TFO12; reverse binding of TFO12 would disrupt the consecutive formation of base triples and therefore reduce the stability of the RNA triplex construct. The TFO12-bound ST-DPX is called ”Short-Tetraloop Triplex” (ST-TPX). The ”Long-Tetraloop Duplex” (LT- DPX) was designed by duplicating the TFO12 binding domain in the duplex stem: 5’- GGG CUA GAG AGA GAA AGA GAG AGA GAA AGU UUC GAC UUU CUC UCU CUC UUU CUC UCU CUA GCC C. The long and asymmetric TFO binding domain can be recognized by TFO24, forming 24 consecutive base triples, and converting the duplex into the ”Long- Tetraloop Triplex” (LT-TPX). The schematic secondary structures of ST-DPX, ST-TPX, LT-DPX and LT-TPX are shown in Fig. 1(a) with bound TFO12 and TFO24 via Hoogsteen hydrogen bonds in the RNA major groove. Note that the cytosine bases in the TFOs have to be protonated (at acidic pH) to form stable C-G·C^+^ base triples.

### RNA Triple Helix Sample Preparation

The double-stranded DNA templates of ST-DPX and LT-DPX were designed with the following sequences respectively: GCC GCC AGT GAA TTC TAA TAC GAC TCA CTA TAG GGC TAG AGA GAG AAA GTT TCG ACT TTC TCT CTC TAG CCC and GCC GCC AGT GAA TTC TAA TAC GAC TCA CTA TAG GGC TAG AGA GAG AAA GAG AGA GAG AAA GTT TCG ACT TTC TCT CTC TCT TTC TCT CTC TAG CCC and purchased from IDT (Coralville, IA). Each 1.0 *µ*g of DNA template was mixed with 10 *µ*L of RiboMAX Express T7 2X Buffer, 2 *µ*L of T7 Express Enzyme Mix (Promega, Fitchburg, WI), and sufficient RNase-free water to make one-reaction volume of 20 *µ*L. We used four parallel batches, a total of 40 T7 reactions, for both ST-DPX and LT-DPX RNA constructs and the mixtures were kept at 37 °C for 12 hours. We used ethanol precipitation at -80 °C to stop the T7 reaction and to condense the RNA products and impurities. The precipitated RNA pellets were dried, re-dissolved and purified at pH 7.0 using a Mono Q 5/50 GL anion exchange column (GE Healthcare, Chicago, IL). The elutions containing the ST-DPX or LT-DPX were collected, concentrated and buffer exchanged using 10-kDa concentrators into the Hairpin Annealing Buffer (HAB) containing 40 mM NaCl, 20 mM sodium 3-(N-morpholino)propanesulfonic acid (Na-MOPS), 50 *µ*M ethylenediaminetetraacetic acid (EDTA), pH 7.0. The RNA samples were annealed in HAB. A portion of the prepared ST-DPX and LT-DPX samples was buffer exchanged to meet experimental conditions.

The anti-sense single-stranded TFO12 and TFO24 were purchased from IDT (Coralville, IA). We prepared the cold (0 °C) Triplex Annealing Buffer (TAB) containing 200 mM NaCl, 1.0 mM MgCl_2_, 20 mM sodium 2-(N-morpholino)ethanesulfonic acid (Na-MES), 50 *µ*M EDTA, pH 5.5, to reconstitute the TFOs and protonate the cytosine bases. We mixed the DPX and corresponding TFO (ST-DPX + TFO12 or LT-TPX + TFO24) using a 1:1.2 ratio in 0 °C TAB with a total volume of 100 *µ*L. The mixtures were then annealed using a programmable PCR machine. The annealed RNA constructs: ST-TPX and LT-TPX were purified at pH 5.5 using a Mono Q 5/50 GL anion exchange column (GE Healthcare, Chicago, IL). The purified RNA triplexes were further concentrated, and buffer exchanged to either experimental conditions: 200 mM NaCl or 5 mM MgCl_2_ in 20 mM Na-MES, 50 *µ*M EDTA, pH 5.5. Final ST-DPX and LT-DPX concentrations ranged from 400 to 800 *µ*M while STTPX and LT-TPX samples were about 150 to 320 *µ*M. The samples were further diluted at the beamline prior to the solution X-ray scattering experiments.

### RNA Triple Helix Sample Characterization

We performed anion exchange and circular dichroism (CD) experiments to characterize all the RNA samples: ST-DPX, ST-TPX, LT-DPX and LT-TPX.

#### Anion exchange analysis

The anion exchange experiments were conducted using the UV-absorption-coupled AKTA system and the Mono Q 5/50 GL anion exchange column (GE Healthcare, Chicago, IL) at 4 °C. We used 100 mM and 1.0 M NaCl in pH 5.5 as the starting and elution buffers. About 100 *µ*L of RNA samples were manually loaded and injected into the column after equilibration of the system. The fraction of the elution buffer was linearly increased from 0% to 100% in 90 minutes. Conductivity, UV absorption and elution volume were monitored and recorded by the UNICORN software. The UV absorption traces are shown in Fig. S1 as a function of conductivity in mS/cm. In Fig. S1(a), we distinguish ST-TPX from ST-DPX by their different elution conductivities, confirming the formation of ST-TPX. In Fig. S1(b), we demonstrate the sequence-specific binding of the TFO12 by annealing ST-DPX with TFO12 and a 12-nt poly(rU). The poly(rU)_12_ does not bind to ST-DPX and the sample shows both ST-DPX and the poly(rU)_12_ peaks (green). We further annealed LT-DPX with TFO12 and TFO24 separately and the anion exchange traces are shown in Fig. S1(c). Since LT-DPX has two identical TFO12 binding domains, it can bind with one TFO12 or two TFO12s, forming two additional peaks at 50 and 52.5 mS/cm respectively. When annealed with TFO24, LT- TPX was formed and its peak coincides with two-TFO12-bound LT-DPX. Therefore, Fig. S1 shows the sequence-specific binding of the TFOs and therefore the formation of ST-TPX and LT-TPX. Single peak from ST-TPX implies that the ST-DPX has only one TFO12-binding domain and therefore, that ST-DPX is a hairpin monomer. Note that the ST-DPX (blue) and LT-DPX (red) in Fig. S1(b,c) might contain impurities from the T7 reactions.

#### Circular dichroism experiment

The CD experiments were performed on the MOS-500 spectropolarimeter (BioLogic, Seyssinet-Pariset, France) using a wavelength scan from 210 to 320 nm at room temperature. About 20 *µ*L solution sample with nucleic acid concentration of 1.5 *µ*g/*µ*L was loaded into a 0.1-mm UV Quartz U-shaped circular dichroism cuvette. Ten CD traces of the sample and buffer were taken and averaged. The final CD traces were acquired by subtracting the buffer trace from the sample trace. They are shown in Fig. S2. The duplex to triplex transition corresponds to a major shift in wavelength of the 260-nm peak and a dip in the curve at about 230 nm.

#### UV/Vis melting

We further performed UV/Vis melting experiments on the Cary 50 Spectrophotometers (Agilent, Santa Clara, CA) to characterize the ST-DPX and ST-TPX RNA motifs. We loaded 100 *µ*L solution sample in the UV Quartz cuvette and sealed the cuvette to avoid evaporation. The UV absorption at 260 nm was recorded with a linear temperature ramp from 20 to 95 °C over a period of one hour. The raw melting curves and the first derivatives, *dA/dT* , are shown in Fig. S3 for ST-DPX and ST-TPX. The first *dA/dT* peak of ST-TPX at about 45 °C (cyan) corresponds to the dissociation of TFO12 while the second peak at about 85 °C is due to the denaturing of the RNA secondary structure (hairpin) in both ST-DPX and ST-TPX.

## General MD simulation set up

The built structures were immersed in a triclinic cell by extending the simulation box by about 15 A^\x{FFFF}^ in each direction from the RNA surface, giving rise to the box sizes of 7.0x7.0x15.0 *nm*^3^ for LT-TPX/DPX systems and 7.0x7.0x10.0 *nm*^3^ for ST-TPX/DPX. The RNAs were solvated with water and ions to match experimental conditions, see Table S1 for further details.

All simulations were carried out using GROMACS 5.0.5^47^ suite of programs. We computed the long-range interactions with a distance cutoff of 0.12 nm. Van der Waals interactions include dispersion correction, while the electrostatics were treated by particle mesh Ewald^70^ (PME) summation method. The PME scheme used a grid spacing of 0.12 nm with an interpolation of order 4. The covalent bond lengths of the water and nucleic acids were constrained by SETTLE^71^ and LINCS^72^ algorithms, respectively. All simulations employed a time step of 2fs.

The structural search was started with energy minimization, to remove possible bad contacts due to random placement of water molecules and ions. To prepare the structures for Sample and Select, and subsequently for WAXS-driven MD simulations, we employed a series of equilibriation steps that include 2 ns constrained MD at constant pressure and temperature to adjust the volume. Subsequently, we conduct 200 ns constrained MD where heavy atoms of RNA were restrained while water and ions allowed to move. This step is followed by 300 ns unrestrained simulations to sample RNA conformations at canonical ensemble.

### WAXS Computing from MD trajectory

The conformations sampled by simulations were used to compute the WAXS profiles. From each solute system (RNA+ion+water), we constructed a molecular envelope that extends to bulk. We used a cutoff of 12 Å to construct this envelope. In addition to the solute simulations, we ran 20-ns-long MD simulations of buffer environment using NVT ensemble. A buffer simulation contains the ion pairs and water molecules in a periodic box with dimensions and concentration matching the solute system. The bulk simulations allowed accurately treating the solvation shell and the excluded volume effect needed for buffer subtractions. Similar to the solute systems, we used an envelope that extends to 12 Å for the bulk systems. The simulated WAXS curves are obtained by subtracting the buffer intensity from the scattering profile of the solute. Here, the instantaneous intensity at time *t* was obtained by

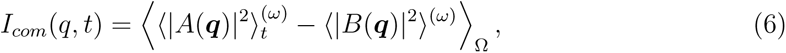

where *A*(***q***) and *B*(***q***) indicate the Fourier transform of the real-time electron density of RNA+water+ion and water+ion systems respectively. 〈·〉_Ω_ and 〈·〉^(^*^ω^*^)^ denote the orientational average and the conformational average at a fixed orientation (*ω*).

### Coupling experimental data with MD simulations

The refinement of the conformational ensemble was achieved by the introduction of an additional energy term *E_hybrid_* = *E_FF_* + *E_W AXS_* to the Hamiltonian^40^. Two different functional forms of the coupling potential *E_W AXS_* were employed: i) non-weighted coupling (Eq. 2)

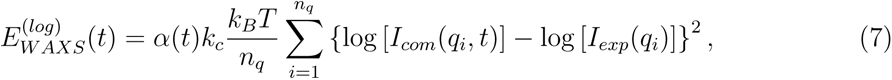

followed by ii) uncertainty-weighted coupling (Eq. 8).

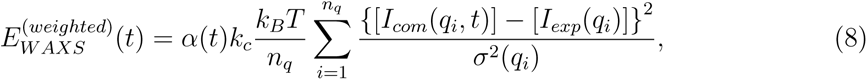

where 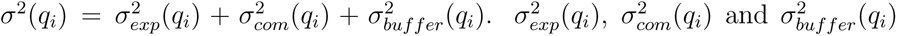 are experimental error, statistical error from computed curves, and systematic error given by the uncertainty of buffer density, respectively. [*I_com_*(*q_i_, t*)] is the real-time WAXS curve from the simulation, while *n_q_* is number of intensity points spanning the specific range of scattering vector *q*. The coefficient *k_c_* here is a constant that adjusts the weight of the WAXS potential *E_W AXS_* compared to the force field *E_FF_* term. The parameter ff(*t*) is a time-dependent function that allows a gradual introduction of the coupling potential at the start of the simulation.

The coupling of the WAXS data to the simulations is accomplished by applying an additional force (Eq. 9) to atomic coordinates:

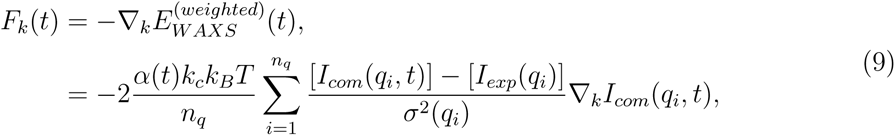

where *r_k_* indicates the gradient with regards to the atomic position of atom *k*. Here, we applied forces only to the solute atoms, i.e. the nucleic acid helices.

### WAXS-driven simulation set up

For WAXS data-driven MD,^53^ the equations of motion were integrated using stochastic dynamics (SD) integrator. We removed the center of mass motion, while we kept other settings remain unchanged from the previous section of **General MD simulation set up**. The scattering amplitudes are averaged by giving more weight to recent conformations using an exponential decaying memory function ^40^. The memory time *τ* was set to 250 ps and enabled the convergence of buffer subtraction over the time interval covering the fluctuations of solvent and RNA backbone. A time interval *t* of 20 ns was used for the switch function *α*(*t*) to avoid strong WAXS-derived bias and to allow sufficient convergence of *I_com_*(*q, t*) before applying the WAXS derived forces.

The computed WAXS curves were reported every 5 ps to monitor progress. The uncertainty of solvent density was set to 0.5%. We used 1500 *q*-vectors for orientational average. Due to the relatively high uncertainty in the far WAXS regime, the structures were refined using the experimental data up to *q* = 10.0 *nm^-^*^1^. The number of restrained *q* points *n_q_* was roughly determined according to the Shannon information theory.^73^ We used *n_q_* =40 for ST-TPX/DPX systems and 50 for LT-TPX/DPX. For non-weighted coupling potential, we used a coupling strength of *k_c_*=100. During uncertainty-weighted coupling, we choose *k_c_*=0.5 for LT-TPX/DPX and *k_c_*=0.4 for ST-TPX/ST-DPX respectively.

The convergence of WAXS-driven MD was evaluated by customized x^2^ metric (Eq. 1) as well as in-house assessment (in section **Calculation of Solution X-ray Scattering Profiles**) by re-computing WAXS profiles from the selected x^2^-converged conformations. Sorted by x^2^ metric, the conformations giving the best agreement with experiments build the conformational ensemble. The data is clustered using the gromos algorithm. The cluster centers were used to represent the corresponding ensembles.

### Residue-residue Contact Analysis

For each residue pair (*ij*) of RNA we compute the contact frequency that posses a specified minimum distance **d** range. Note that **d***_Loop-Duplex_* is measured from the minimum distance between each loop residue and from the closest stem base pair, while **d***_TF O-Duplex_* is estimated from the minimum distance between each TFO residue and its corresponding complementary sequence on the stem.

---

**Table S1:**
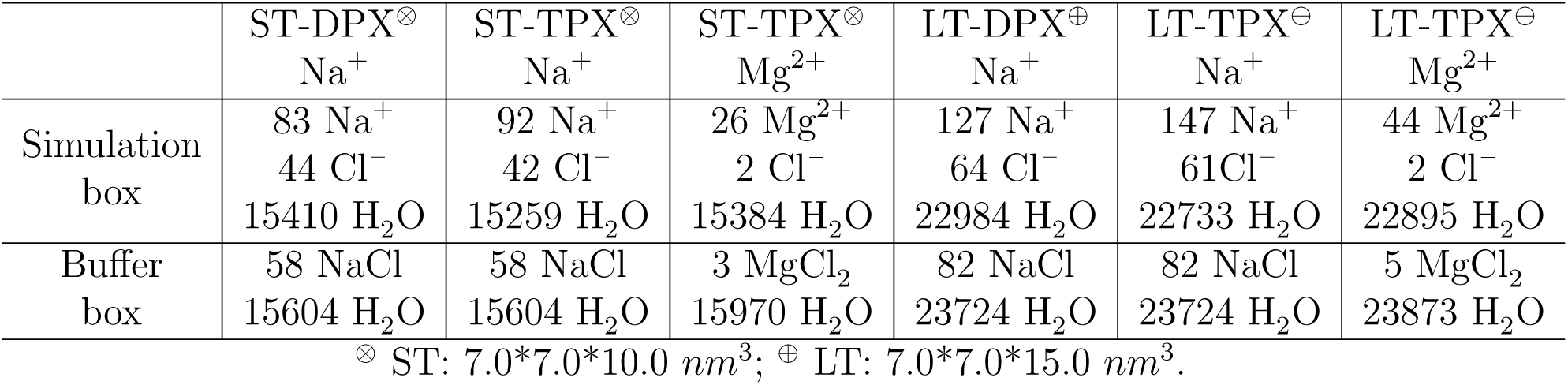
Simulation setup details

**Table S2:** SASBDB access codes of all the experimental solution X-ray scattering profiles used in this work.

|  | ST-DPX | ST-TPX | LT-DPX | LT-TPX |
| --- | --- | --- | --- | --- |
| 200 mM NaCl | SASDKP5 | SASDKS5 | SASDKV5 | SASDKY5 |
| 5 mM MgCl<sub>2</sub> | SASDKR5 | SASDKU5 | SASDKX5 | SASDK26 |

**Figure S1:**
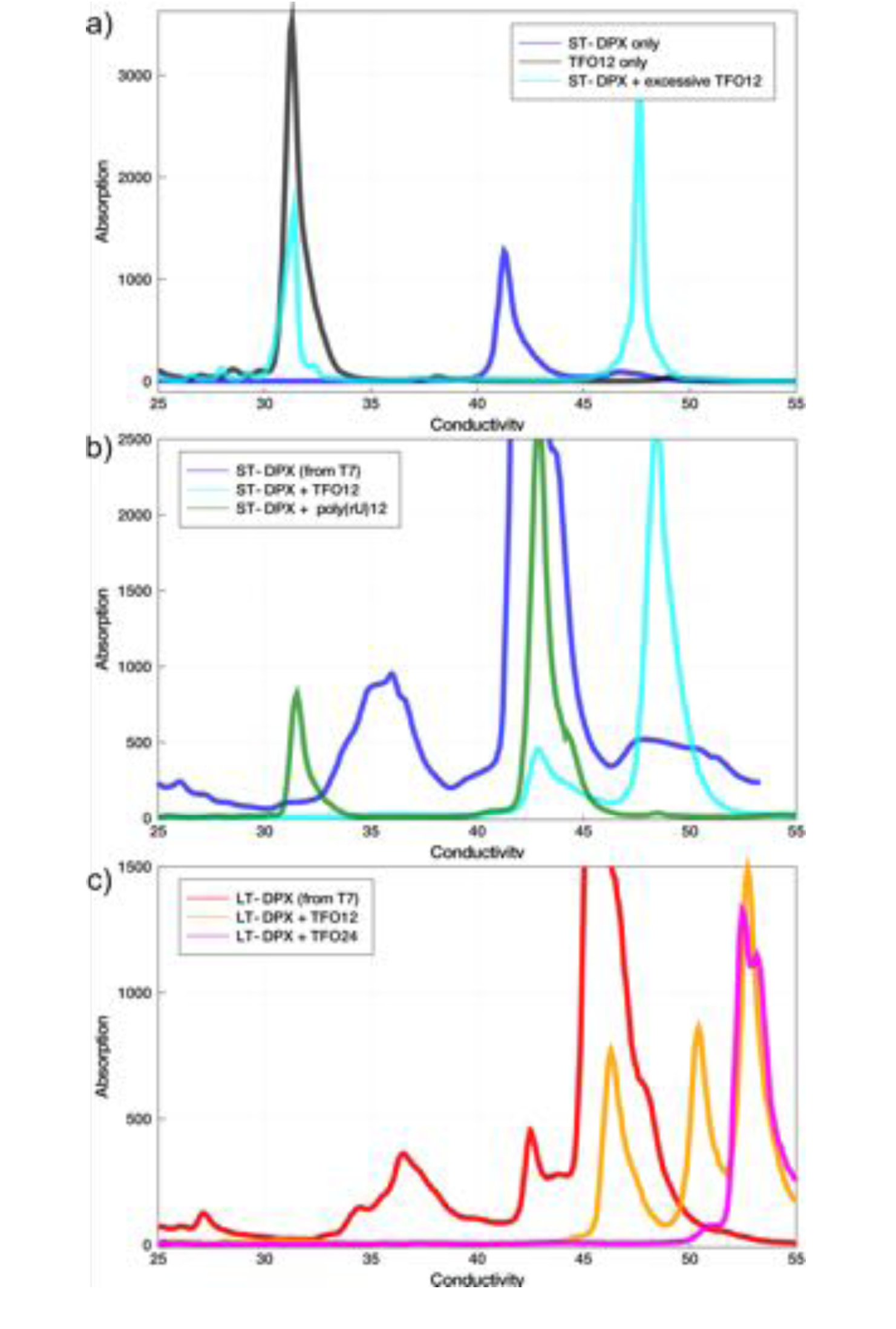
The anion exchange traces (absorption versus conductivity in mS/cm) from triplex characterization experiments. (a) The traces of pure ST-DPX (blue), TFO12 (black) and annealed ST-DPX + TFO12 (cyan). The later-eluted sample at 17.5 mS/cm is the ST-TPX. (b) The traces of ST-TPX (blue), ST-DPX + TFO12 (cyan) and ST-DPX + poly(rU)_12_ (green). The latter does not show binding of the poly(rU)_12_, implying sequence-specific binding of the TFO12. (c) The traces of LT-DPX (red), LT-DPX + TFO12 (orange) and LT-DPX + TFO24 (magenta). The LT-DPX can bind either one or two TFO12s and shows two additional peak at 50.0 and 52.5 mS/cm. The LT-TPX coincides with the latest-eluted sample. Note that ST-DPX (blue) and LT-DPX (red) in (b-c) contains impurities from T7 reactions.

**Figure S2:**
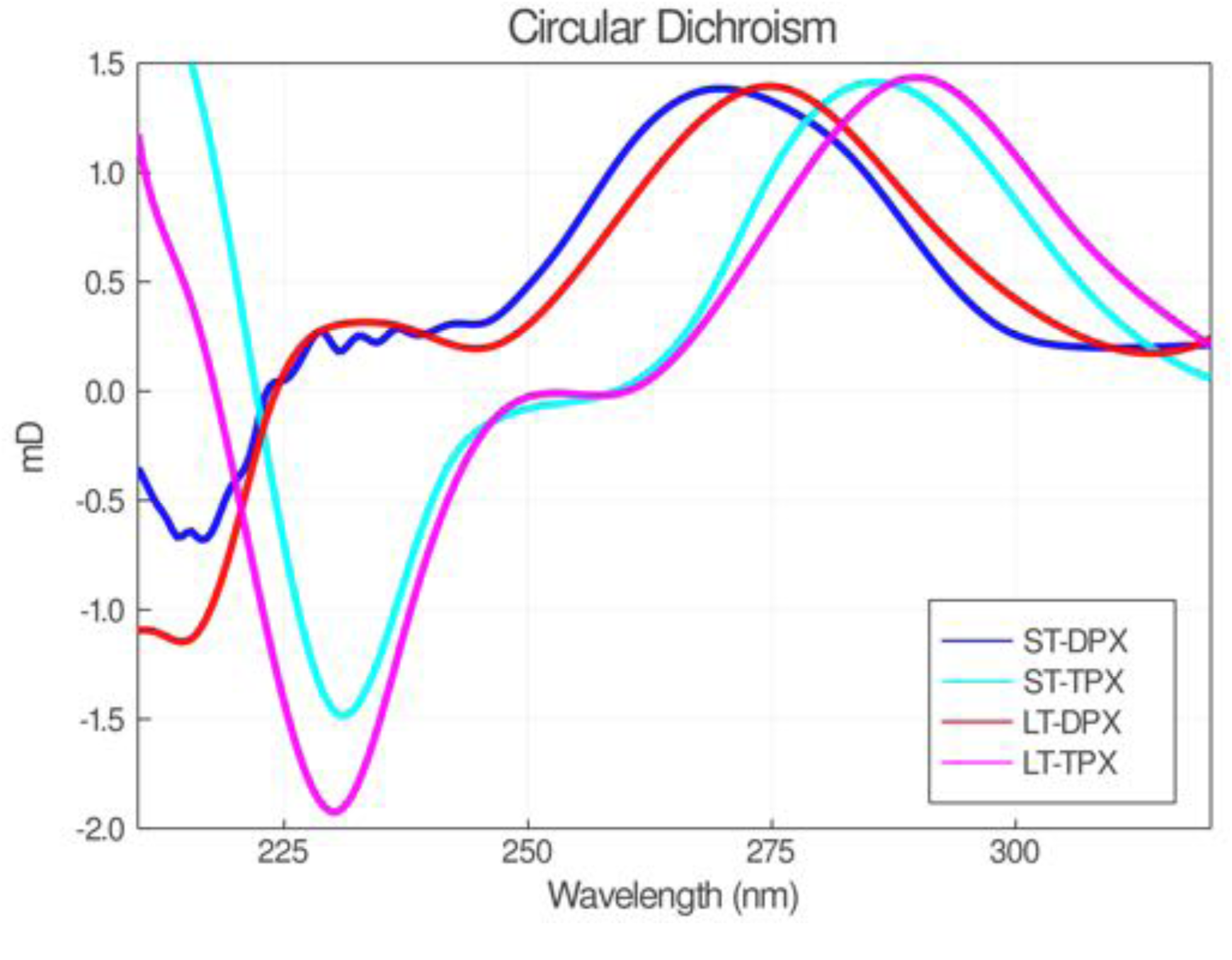
The concentration-normalized circular dichroism (CD) traces of ST-DPX, LT- DPX and their corresponding triplexes: ST-TPX and LT-TPX. The y-axis is in millidegree for 1.5 *µ*g/*µ*L sample in 0.1-mm-thick cuvette. The CD spectra reflect more of the 3D structures of the RNA than the individual bases. The transition from RNA duplex to RNA triplex is reflected by the shift of the main peak from *⇠* 260 nm to longer wavelength.

**Figure S3:**
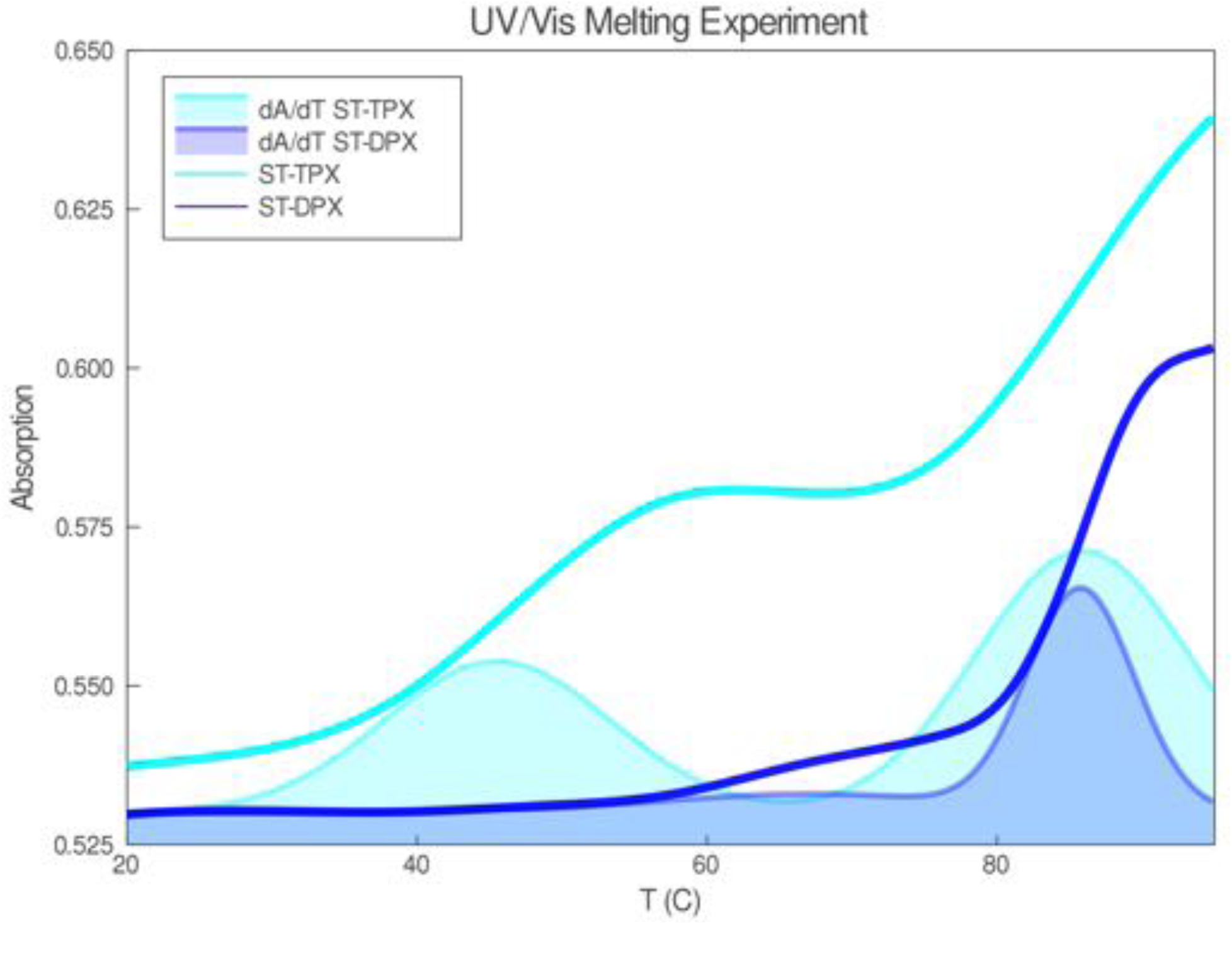
The UV/Vis melting curves are shown as absorption (A) versus temperature (T) in °C for ST-DPX (blue) and ST-TPX (cyan). The first derivatives, *dA/dT* , are also shown. The first change (increasing slope) of ST-TPX at about 45 °C is the dissociation of the TFO12 while the increase at about 85 °C corresponds to the denaturing of the RNA hairpin secondary structure in both ST-DPX and ST-TPX motifs.

**Figure S4:**
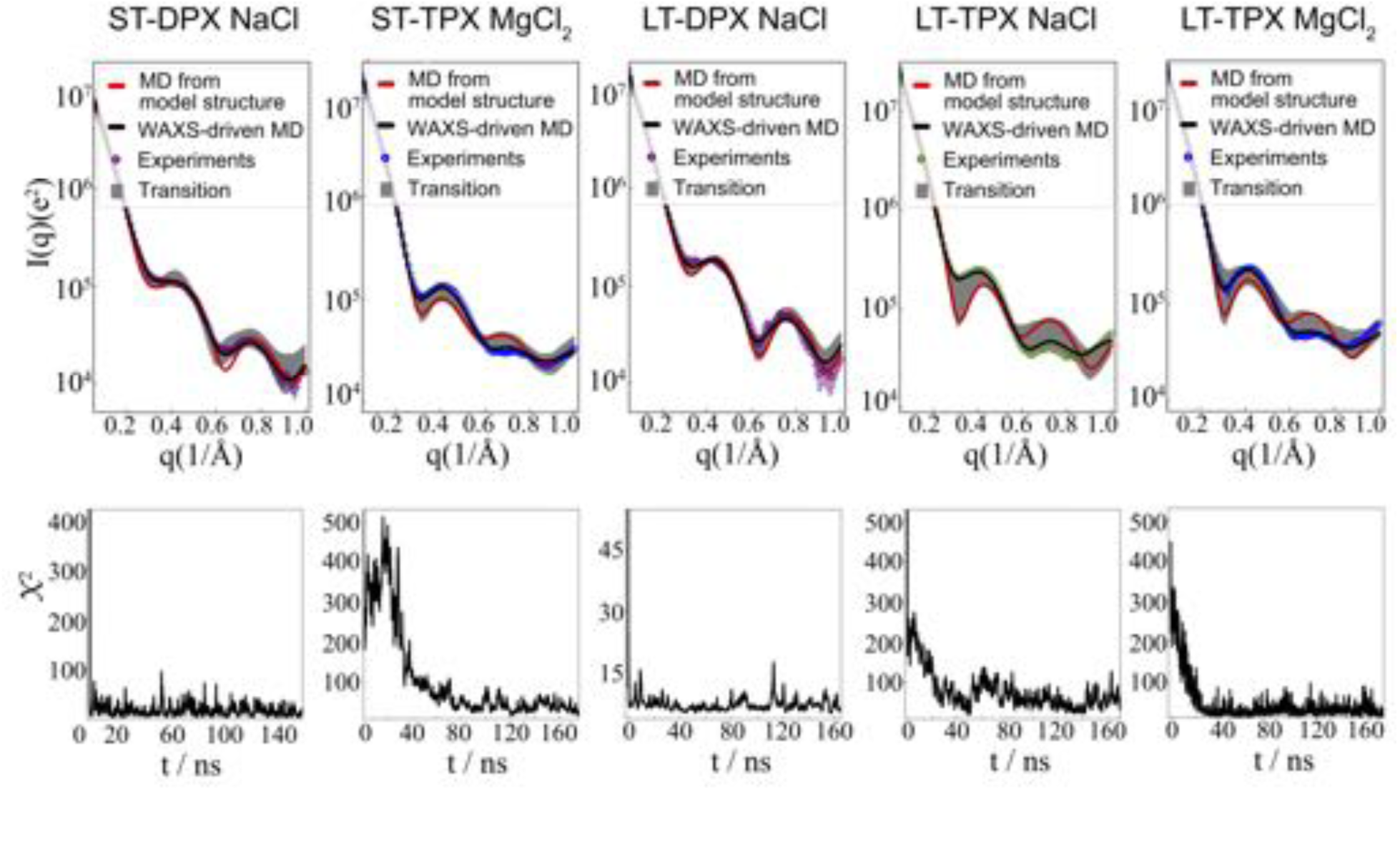
The transition of WAXS profile computed from simulation and variation of customized metric x^2^ during WAXS-driven MD.

**Figure S5:**
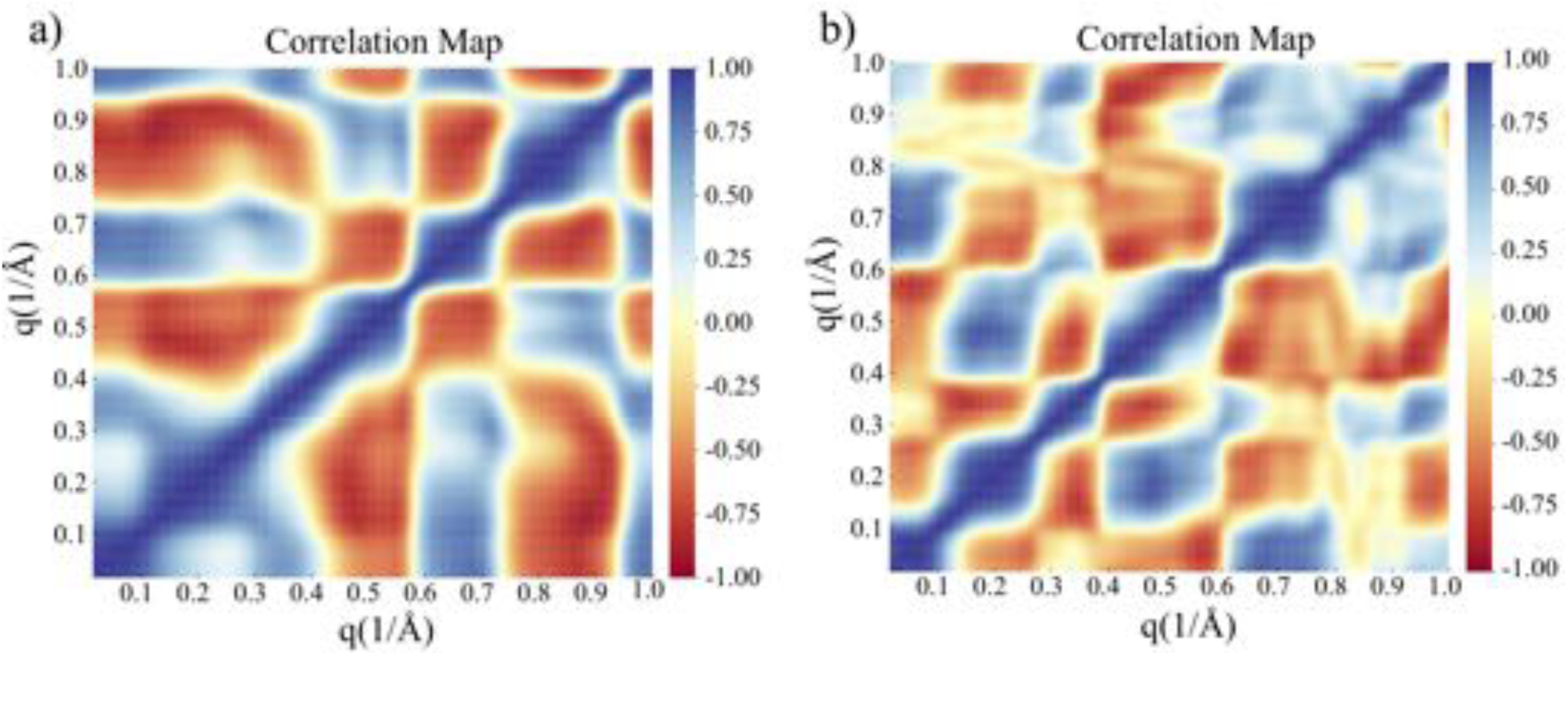
The correlation map among normalized deviations at different *q* in Eq. 4 (c*_k_* (*q_i_*)) for (a) ST-TPX and (b) LT-TPX.

**Figure S6:**
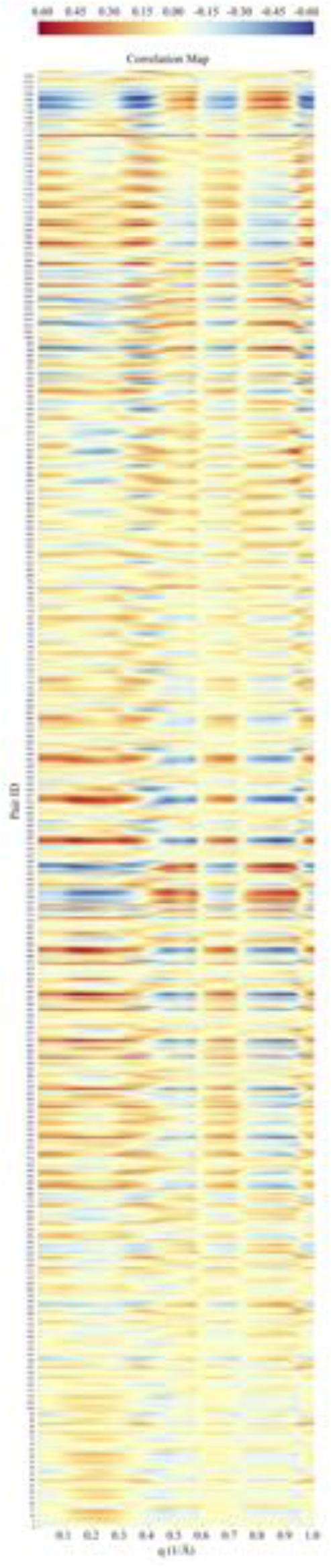
Full correlation map between normalized deviations of pairwise distance and WAXS at different *q* in Eq. 4 for ST-TPX. The sequential numbers (bottom-up) of y axis locate the residue pairs, in the order of r1-r2,r1-r3,…,r1-r52,r2-r3,…,r51-r52.

**Figure S7:**
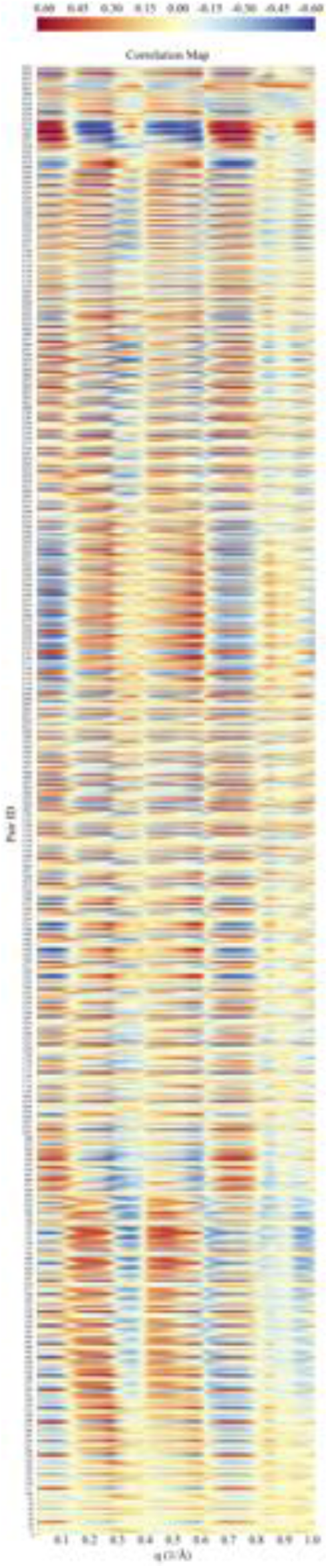
Full correlation map between normalized deviations of pairwise distance and WAXS at different *q* in Eq. 4 for LT-TPX. The sequential numbers (bottom-up) of y axis locate the residue pairs, in the order of r1-r2,r1-r3,…,r1-r88,r2-r3,…,r87-r88.

**Figure S8:**
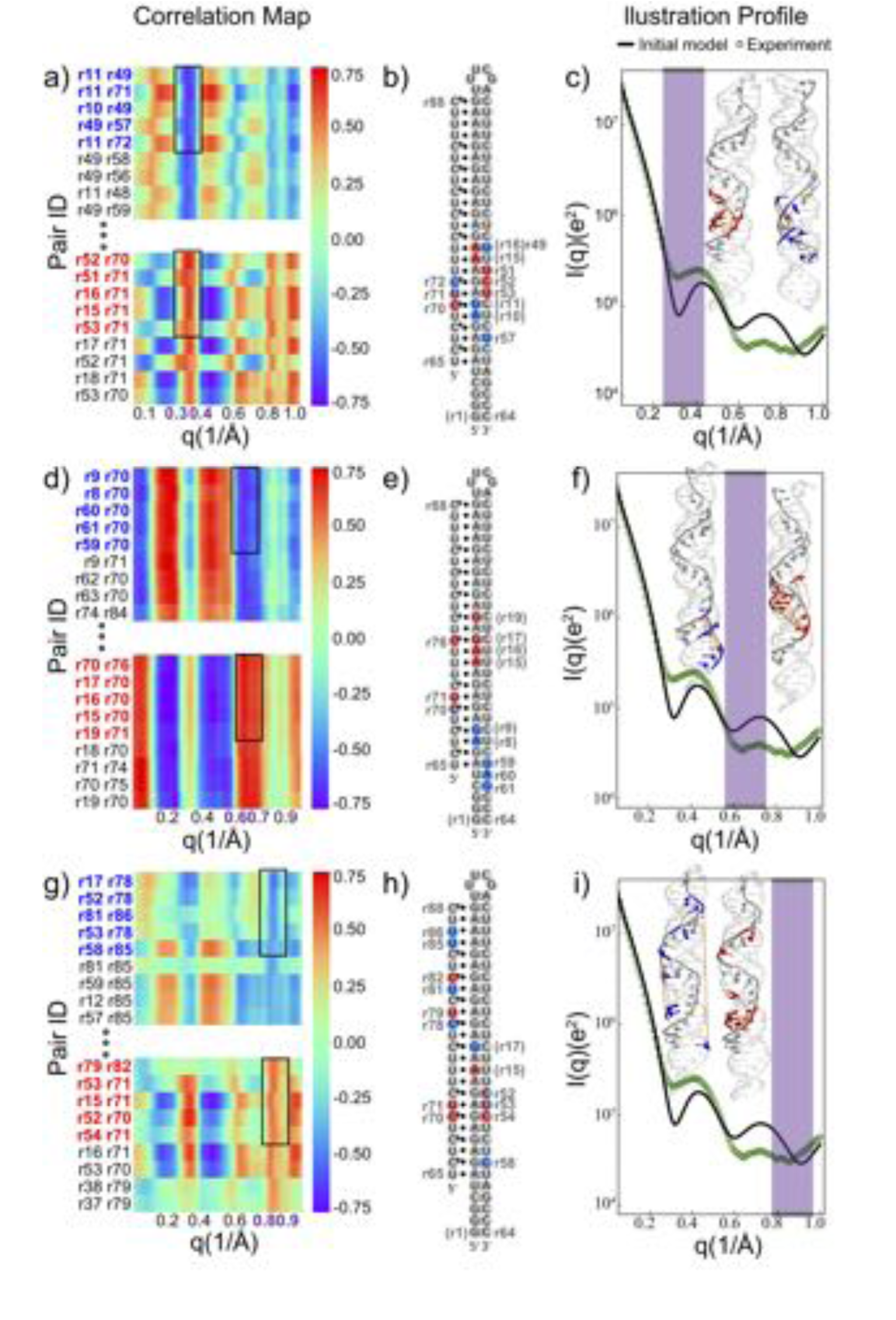
Correlation analysis based on standard MD of LT-TPX in 200 mM NaCl, connecting residue-residue pairwise distance and normalized WAXS deviation between simulation and experiment. See detailed description in Fig. 4 and main text.

